# Sailing in rough waters: examining volatility of fMRI noise

**DOI:** 10.1101/2020.06.19.161570

**Authors:** Jenni Leppanen, Henry Stone, David J. Lythgoe, Steven Williams, Blanka Horvath

## Abstract

**Background:** The assumption that functional magnetic resonance imaging (fMRI) noise has constant volatility has recently been challenged by studies examining heteroscedasticity arising from head motion and physiological noise. The present study builds on this work using latest methods from the field of financial mathematics to model fMRI noise volatility.

**Methods:** Multi-echo nd human fMRI scans were used and realised volatility was estimated. The Hurst parameter *H* ∈ (0, 1), which governs the roughness/irregularity of realised volatility time series, was estimated. Calibration of *H* was performed pathwise, using well-established neural network calibration tools.

**Results:** In all experiments the volatility calibrated to values within the rough case, *H* < 0.5, and on average fMRI noise was very rough with 0.03 < *H* < 0.05. Some edge effects were also observed, whereby *H* was larger near the edges of the phantoms.

**Discussion:** The findings suggest that fMRI volatility is not only non-constant, but also substantially more irregular than a standard Brownian motion. Thus, further research is needed to examine the impact such pronounced oscillations in the volatility of fMRI noise have on data analyses.

## 1. Introduction

A given functional magnetic resonance imaging (fMRI) blood oxygenation level dependent (BOLD) time series can be defined as

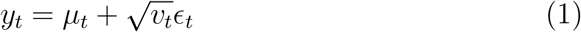

where the *μ_t_* is the mean, *ϵ_t_* is a one dimensional noise process, and *v_t_* is the volatility of the noise process. Detrending is typically conducted as part of preprocessing to remove signal drift. Thus, a given fMRI time series is often assumed to be a constant process, indicating that *v_t_* in equation (1) could be replaced by constant *v*. This assumption, however, has recently been challenged and there has been increasing interest in exploring time-dependent properties of fMRI noise [1, 2, 3, 4, 5, 6]. It has been shown that factors such as head motion and physiological processes including respiration and pulse can introduce heteroscedasticity to the time series [1, 2, 3, 7]. Heteroscedasticity in turn has been found to complicate the linear modelling, which has led to the introduction of several statistical models to counteract the impact of these artefacts [1, 2, 7]. One limitation of these models is that they cannot explain non-constant volatility arising from unknown or uncontrollable sources, such as scanner noise.

As volatility of a time series cannot be directly observed, a plethora of deterministic and stochastic models have been proposed to estimate it in financial returns data [8, 9, 10]. Over the years, direct comparisons of different volatility models have shown that stochastic models, which assume that logarithm of the volatility process behaves like standard Brownian noise with Hurst parameter *H* = 0.5, outperform their deterministic, data-driven counterparts providing a better fit to data [11, 12, 13]. This assumption implies in particular that volatility is not constant^1^, and exhibits an oscillatory behaviour on any finite time interval. This oscillatory behaviour is governed by a parameter *H* which in the Brownian case takes the value *H* = 0.5.

More recently rough stochastic volatility models have been considered (see [14, 15, 16, 17, 18]) where the parameter *H* is allowed to vary in the range *H* ∈ (0, 1). In these models, as mentioned above, the parameter *H* governs the oscillations of the volatility process; the lower the parameter *H*, the stronger the oscillations on any finite interval. In particular, the values *H* ∈ (0, 0.5) correspond to the *rough* case (i.e. rougher paths than a standard Brownian motion). Figure 1 shows the roughness/irregularity of volatility paths for different *H* values. As *H* approaches 0 the paths become more irregular/rough.

**Figure 1:**
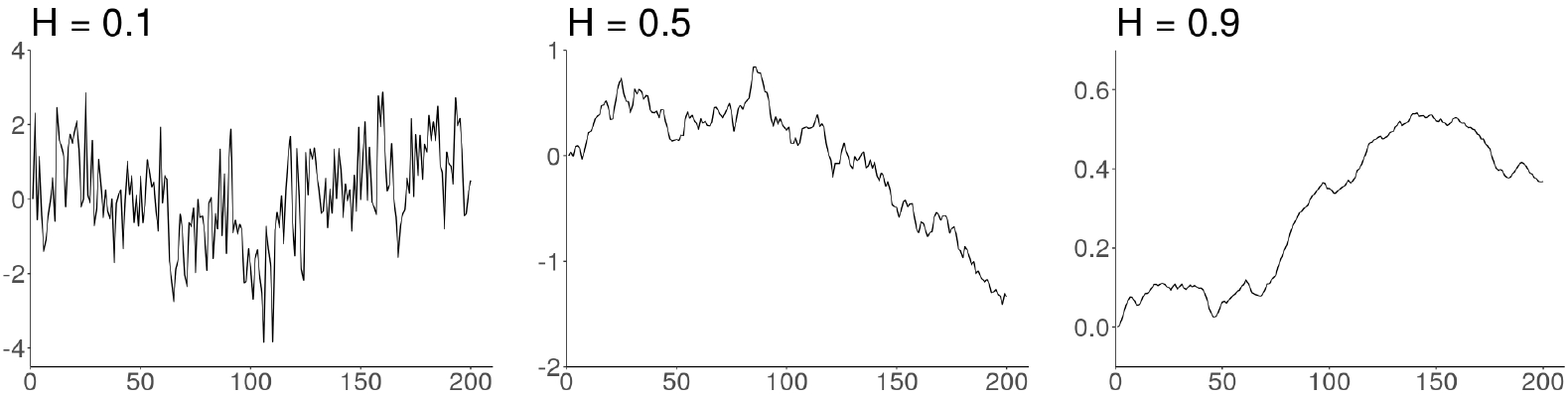
Three sample paths of fractional Brownian motion with *H* = 0.1, *H* = 0.5, and *H* = 0.9

A rough stochastic volatility model, the rough Bergomi (rBergomi) model introduced in [15] by Bayer Friz and Gatheral, is described by the system

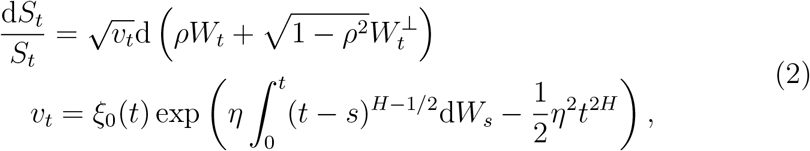

where *W* and *W* ^⊥^ represent two independent standard Brownian motions with *ρ* ∈ [−1, 1], *η* > 0 describes the volatility of volatility, and *ξ*_0_(·) describes the initial variance curve, which we assume to be constant. Our motivation for choosing the mean reverting, driftless rough Bergomi model (2) derives from the practice of *detrending* mentioned above. As it is common practice to remove linear drifts from fMRI prior to further analysis, such a driftless model would be a good fit to the data. Volatility processes simulated using the rBergomi model exhibit remarkable similarity to realised volatility processes [18, 15]. Furthermore, the rBergomi and other rough models introduced since have been found to provide important improvements to forecasting volatility [15, 19, 9, 10].

In addition to improving forecasting accuracy, rough models can be used to assess smoothness of a given process by estimating the parameter *H* [17, 18, 20, 21]. Estimating the parameter *H* can provide information about the extent of heteroscedasticity in the series, but requires access to the realised, historical volatility process, which cannot be directly observed. To bypass this difficulty, in finance intra-day data such as 5-minute asset prices returns are used to estimate daily realised volatility [22, 23, 24]. The daily estimates are then combined to form a realised volatility process, providing information about daily variances in an asset price over the course of months or years.

Considering recent calls to explore the possibility of implementing models from the field of financial mathematics to fMRI [6, 25] and the visual similarities between financial returns data and fMRI BOLD signal (Figure 2), such an approach could be applied to fMRI data as well to examine time-dependent behaviour in volatility of the noise process. Utilising multi-echo acquisition, the data from each echo could be used as intra-time point data to estimate volatility. Thus, in a manner similar to standard combination of data from each echo time, we can produce a realised volatility series. These series could then be used to estimate the smoothness of volatility in fMRI data using models such as the rBergomi.

**Figure 2:**
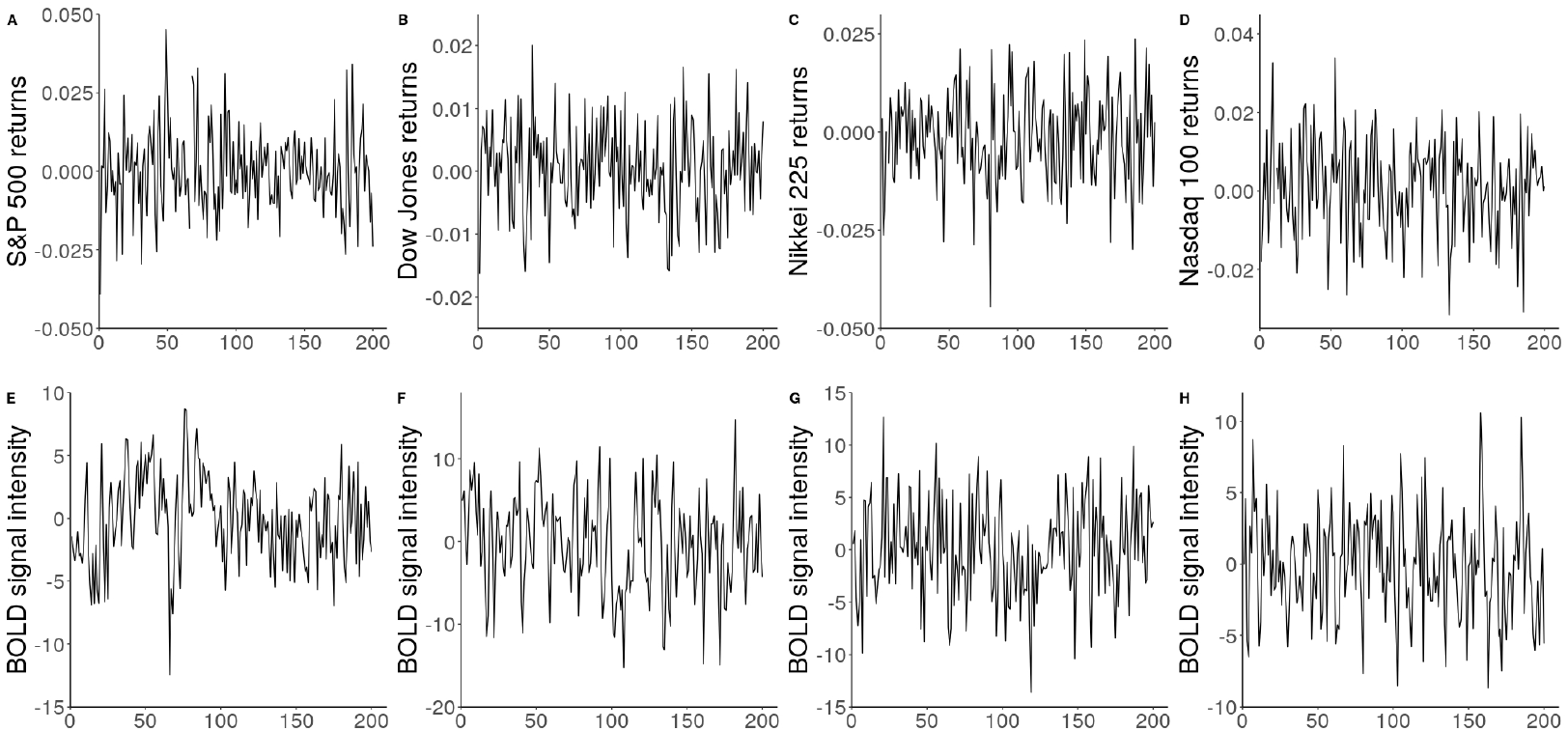
Financial returns and fMRI BOLD series A. S&P 500 returns over 200 days during the year 2000; B. Dow Jones returns over 200 days over the year 2004; C. Nasdaq 225 returns over 200 days during the year 2003; D. Nikkei 100 returns over 200 days during the year 2000; E. Demeaned BOLD signal from voxel [28,14,26]; F. Demeaned BOLD signal from voxel [12,27,30]; G. Demeaned BOLD signal from voxel [17,36,24]; H. Demeaned BOLD signal from voxel [37,19,15]. All returns data is from the Oxford-Man database https://realized.oxford-man.ox.ac.uk/. All fMRI data is from participant sub-17821 from dataset ds000258 available at https://openneuro.org/datasets/ds000258.

Estimating rBergomi model parameters, including *H*, is computationally expensive and often relies on the use of Monte Carlo based calibration methods [26, 27]. This limits the use of this model in practice despite the benefits it offers [18, 28, 15]. Recently, neural networks have been proposed as an efficient way to solve the calibration problem [29, 30, 31, 32]. Neural networks provide a powerful way of identifying relationships between input parameters and model output and can be particularly useful for models that do not have closed-form solution [29, 30]. Recent work found that neural network calibration framework can be successfully applied to a range of rough stochastic volatility models to aid accurate pricing and hedging [29, 33]

The aim of this paper was to conduct an exploratory empirical study examining the volatility of fMRI noise. We were specifically interested in exploring whether volatility of fMRI noise exhibits time-dependent behaviour that cannot be explained by factors such as head motion and physiological noise. We aimed to collect multi-echo fMRI signal from a phantom to examine thermal noise. We also aimed to examine whether volatility patterns observed in the phantom data were present in noise in human scans. To achieve this aim, multi-echo resting state data was extracted from the ventricles of four participants from two different datasets.

Observations collected at each echo time were treated as intra-time point data and were used to estimate realised volatility. The roughness of the realised volatility was assessed by estimating the Hurst parameter *H*, which was accomplished with using neural network calibration tools. As the study was exploratory in nature we did not have prior hypotheses. However, considering the visual similarities between many financial returns and fMRI BOLD series, we anticipated that the estimated *H* of the realised volatility processes was in the rough volatility range, 0 < *H* < 0.5.

## 2. Material and methods

### 2.1. FMRI data acquisition

#### Phantom data

Two MRI phantoms filled with liquid material was used to acquire multi-echo fMRI signal consisting entirely of thermal noise. The data were acquired with two different 3 Tesla GE Discovery MR750 units using 32-channel receive only head coils (Nova Medical, Wilmington, MA, USA). This was done to ensure the findings were not unique to a specific scanner. The functional multi-echo echo planar imaging (EPI) data consisted of 200 volumes and each volume consisted of 18 slices with the following parameters: 2.5 second repetition time (TR), 80 flip angle, 64 × 64 acquisition matrix, 3 mm slice thickness with 4 mm slice gap. The fMRI slices were acquired in an ascending order and eight echo times were used: 12 ms, 28 sm, 44 ms, 60 ms, 76 ms, 92 ms, 108 ms, 124 ms. Eight echo times were used as this was the maximum number of echoes that can be acquired with the MR units used.

#### Human data

Multi-echo resting state data from two different datasets, ds000258 (https://openneuro.org/datasets/ds000258/versions/1.0.0) and ds000210 (https://openneuro.org/datasets/ds000210/versions/00002), were used to examine whether patterns identified in the phantom data could be seen in vivo. The ds000258 data were acquired with a Siemens Trio 3 Tesla MRI scanner using 32-channel receive only head coil. T1-weighted magnetization prepared rapid gradient echo (MPRAGE) sequence was used to acquired the anatomical data with the following parameters: 1 mm slice thickness and 1.1 second inversion time. The functional multi-echo EPI data consisted of 239 volumes and each volume consisted of 32 oblique slices with the following parameters: 2.47 second TR, 78 flip angle, 64 × 64 matrix size, and 4.4 mm slice thickness with 10% slice gap. Alternating slice acquisition was used with ascending interleaved order and four echo times were used: 12 ms, 28 ms, 44 ms, and 60 ms.

The data from the second dataset, ds000210, was acquired with a 3 Tesla GE Discovery MR750 unit using a 32-channel receive only phased-array head coil. T1-weighted MPRAGE sequence was used to acquired the anatomical data with the following parameters: 2530 ms TR, 1 mm slice thickness, and 1.1 second inversion time. The resting state multi-echo EPI data consisted of 204 volumes and each volume consisted of 46 axial slices. The following parameters were used to acquire the data: 3.0 second TR, 83 flip angle, 72×72 matrix size, and 3.0 mm isotropic voxels. The slices were acquired in inferior-superior interleaved order and three echo times were used: 13.7 ms, 30.0 ms, and 47.0 ms.

### 2.2. FMRI data preprocessing

#### Phantom data

The phantom data were preprocessed using SPM12 (http://www.fil.ion.ucl.ac.uk/spm). Each echo was preprocessed separately to ensure the echoes could be used as intra-TR data to estimate realised volatility. The following preprocessing steps were taken: slice timing correction was applied first with the middle slice used as a reference slice. Although no motion was expected, the data were realigned and resliced to correct for head motion and estimate six rigid body transformations. Prior to combining the echoes and estimating realised volatility linear model based de-trending was conducted.

#### Human data

As with the phantom data, SPM12 was used to preprocess the human data one echo at a time to enable estimation of realised volatility. The following preprocessing steps were taken: slice timing correction with the middle slice serving as a reference slice, and realignment with reslicing was used to correct for head motion and estimate six rigid body transformations. The anatomical data were then segmented into grey matter, white matter, cerebrospinal fluid, and skull, after which the anatomical data were co-registered with the mean functional image.

After preprocessing, the six rigid body transformations were used to calculate framewise displacement using the *spmup FD* function (https://github.com/CPernet/spmup/blob/master/QA/spmup_FD.m). Framewise displacement was then used to determine which participants had the least amount of head motion. From each dataset, two participants who moved the least were selected (Supplementary Table 1), the data was subjected to linear model based de-trending, and then taken forward for further analysis. Additionally, to study the noise present in vivo, the anatomical scans were used to create ventricle masks. Studying signal from the ventricles enabled us to examine the volatility of the combination of scanner and physiological noise while avoiding contamination from true brain signal. Thus, only resting state data extracted from the ventricles were used for further analysis to estimate realised volatility and examine its roughness.

### 2.3. 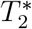-weighted realised volatility

As our data from the phantoms and ventricles contains only noise, we can re-write equation (1) at a given time point *t*…*T* as

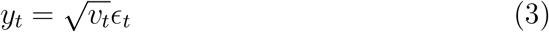

*μ_t_* = 0 as no true brain signal is present.

Observations from each echo time *n* up to the last echo *N* were treated as intra-TR data which were used to estimate realised volatility for each point in the time series. To follow standard procedures and take into consideration the fact that fMRI signal decays rapidly (Supplementary Figures 1-6), the observations from each echo time were weighted to avoid bias [34]. The weighting was based on 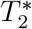 estimates, which were calculated in accordance with methodology used in tedana [35, 36, 37]:

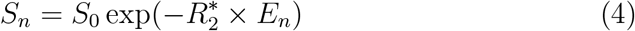

where *S_n_* represents the signal intensity at a given echo time *n*, 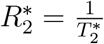, *E* represents the echo time in milliseconds, and *S*_0_ represents the signal intensity at *E* = 0. The value of 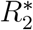 is solved by a log-linear regression.

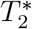-based weights were then calculated as follows

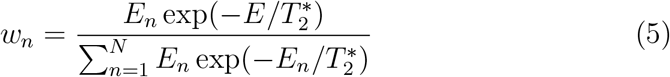

The weights were used to estimate the mean of the fMRI noise processes at each echo time *n* at each time point, *t* = 1…*T*.

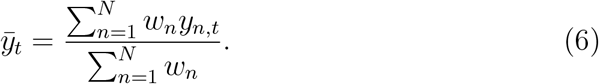

Realised volatility at each time point, *t* = 1…*T*, was estimated by calculating variance between observations at each echo time *n*.

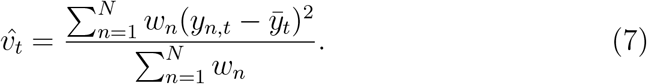

The estimated 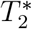-weighted variance, 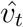, served as a proxy of the unobserved volatility process and was used to investigate the smoothness of the fMRI noise series. As fMRI noise is believed to not exhibit similar exponential decay as true brain signal, we wanted to illustrate that the 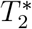-weighting used did not unduly impact the present findings by presenting analyses using non-weighted realised volatility data in the Supplementary Materials. The analyses using non-weighted data produced results which mirror those reported here.

### 2.4. Estimating roughness of realised volatility

We examined the roughness of fMRI noise volatility by adopting a neural network calibration method established in [31]. The rBergomi model was chosen to simulate the training data because it produces driftless, mean reverting processes which closely resemble fMRI data. Roughness of the volatility series was examined by estimating the *H* parameter. In addition to examining the roughness of the volatility paths, we also wanted to extract information about the volatility of volatility by simultaneously estimating the *η* parameter. If any of the fMRI noise volatility series had constant volatility the estimated *η* = 0 and if the volatility was not constant *η* > 0.

#### 2.4.1. Neural network architecture

To estimate the roughness and volatility of volatility of the fMRI noise volatility series we used a one-dimensional feed-forward convolutional neural network (CNN) [31]. This approach has been previously shown to accurately estimate the Hurst parameter *H* and outperform other methods such as the least squares method both in terms of accuracy, as measured using root mean squared error (RMSE), and speed. A further introduction to neural networks is given in Appendix A; very simply one can think of a neural network as a composition of affine and non-linear functions that approximates a mapping of inputs to outputs.

The CNN consisted of three kernel layers with kernel size 20. The first convolutional layer had 32 kernels followed by a dropout layer with dropout rate of 0.25, the second had 64 kernels followed by a dropout layer with dropout rate of 0.25, the third had 128 kernels followed by a dropout layer with dropout rate of 0.4, and the fourth dense layer had 128 units followed by a dropout layer with dropout rate of 0.3. Leaky ReLU activation functions followed each layer with *α* = 0.1 and max pooling layers with size 3 were added between each kernel layer. See [31] for rationale of this architecture and hyperparameter choice.

#### 2.4.2. Neural network training and test

Altogether, 50,000 sample paths of the normalised rBergomi model log-volatility process, 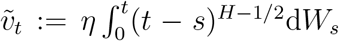, were simulated. For each of the 50,000 sample paths simulated, 200 time points were used and *H* ~ Unif(0, 1.0) and *η* ~ Unif(0, 3.0). Hyperbolic tangent was used to scale *η*. Stone provides a rigorous mathematical justification for this set up [31, Section 3.2, p382]. The sample paths were generated using classical methodology which utilises the Cholesky decomposition to achieve exact distribution of the log-volatility paths (https://github.com/jennileppanen/fmri_vol). The sampling was conducted in a manner that ensured that each sample path had a unique *H* and *η* enabling better fitting to varying fMRI noise log-volatility processes.

We took a nested cross-validation approach whereby the simulated sample paths were first divided into training and test datasets with 30% holdout. The training dataset was then further divided into training and validation sets with 20% hold out. Thus, the training dataset consisted of 28,000 training and 7,000 validation sample paths and the final test dataset included 15,000 sample paths.

### 2.5. Evaluation of CNN H and η parameter

estimation The performance of the trained CNN was assessed by calculating the RMSE between the predicted 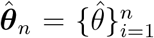 and true 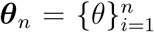 parameters, where 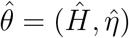 and *θ* = (*H*, *η*).

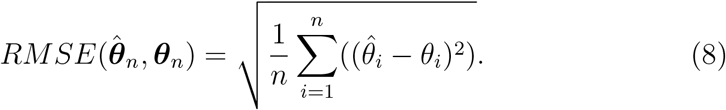

In the present study, the test error was small, *RMSE* = 0.065, and the relationship between predicted and true *H* and *η* in Figure 3 were strongly linear.

**Figure 3:**
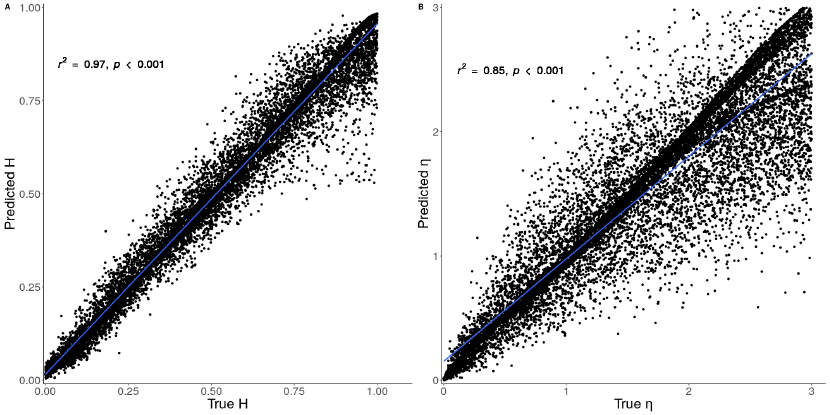
Scatter plot showing the correlation between predicted and true *H* parameter

the parameter *H* governs three aspects of fractional Brownian motion at the same time: the self-similarity, the roughness of the paths (the oscillation) and the autocorrelation of the time series. Therefore, performance of the CNN was additionally evaluated by examining agreement between the estimated *H* parameters and the memory in the fMRI noise log-volatility series. Agreement between the CNN *H* parameter estimates and memory was evaluated by conducting a Spearman correlation test. Memory was estimated by fitting autoregressive fractionally integrated moving average (ARFIMA) [0, *d,* 0] model to the log-volatility data and calculating the *d* parameter:

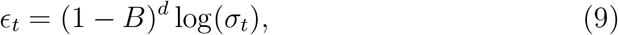

where *B* is the backshift operator and *d* represents the memory parameter to be calculated.

0 < *d* < 0.5 indicates the series is a stationary, mean reverting long memory process, while *d* < 0 indicates the series is anti-persistent short memory process. 0.5 < *d* < 1 indicates the series is a mean reverting, non-stationary long memory process. Although the relationship between smoothness of log-volatility processes and long memory is a complicated one [18, 38, 39, 40], this correlation will give us an indication of the performance of the CNN in estimating *H*.

## 3. Results

### 3.1. Estimated roughness and volatility of volatility

The summary statistics of the estimated *H* parameter of the realised log-volatility series in phantom and human data from the ventricles are presented in Table 1. On average the log-volatility series were rough, but the average *H* parameter estimates were somewhat higher in the data extracted from the ventricles (human) than in the phantom data. This could be because the phantom data should only contain scanner noise while the data extracted from the ventricles should include both scanner and physiological noise. In the phantom data there was also substantial variability in the *H* parameter estimates. The maximum estimated *H* remained under 0.5, suggesting that despite the substantial variability fMRI noise was rough across the phantoms. Similar variability was not observed in the data extracted in the ventricles and the maximum *H* parameter estimates were smaller in the human data. Finally, all *η* > 0 suggesting that there were no fMRI noise volatility processes that had constant volatility.

**Table 1:** Estimated *H* and *η* parameters in phantom and human data

|  |  | Phantom data |  | ds000528 |  | ds000210 |  |
| --- | --- | --- | --- | --- | --- | --- | --- |
|  |  | Phantom 1 | Phantom 2 | sub-17821 | sub-21300 | sub-28 | sub-30 |
| $H$ | Mean | 0.020 | 0.019 | 0.020 | 0.021 | 0.031 | 0.042 |
|  | SD | 0.021 | 0.015 | 0.009 | 0.010 | 0.013 | 0.020 |
|  | Max | 0.498 | 0.396 | 0.042 | 0.052 | 0.107 | 0.127 |
|  | Min | 0.0005 | 0.001 | 0.003 | 0.004 | 0.004 | 0.008 |
| $\eta$ | Mean | 0.232 | 0.234 | 0.331 | 0.345 | 0.462 | 0.526 |
|  | SD | 0.111 | 0.071 | 0.053 | 0.058 | 0.098 | 0.134 |
|  | Max | 4.503 | 3.020 | 0.443 | 0.493 | 1.058 | 1.193 |
|  | Min | 0.036 | 0.055 | 0.203 | 0.184 | 0.235 | 0.285 |

Human data was extracted from the ventricles. SD = standard deviation; ds000258 and ds000210 refer to the two Openneuro. datasets used.

### 3.2. Spatial pattern in estimated Hurst parameters

Figure 4 shows how the estimated *H* varied from region to region across the phantoms and Figures 5 and 6 show *H* parameter estimates in the ventricles. Log-volatility series associate with the maximum, minimum, and *H* parameter estimates close to the mean are also shown.

**Figure 4:**
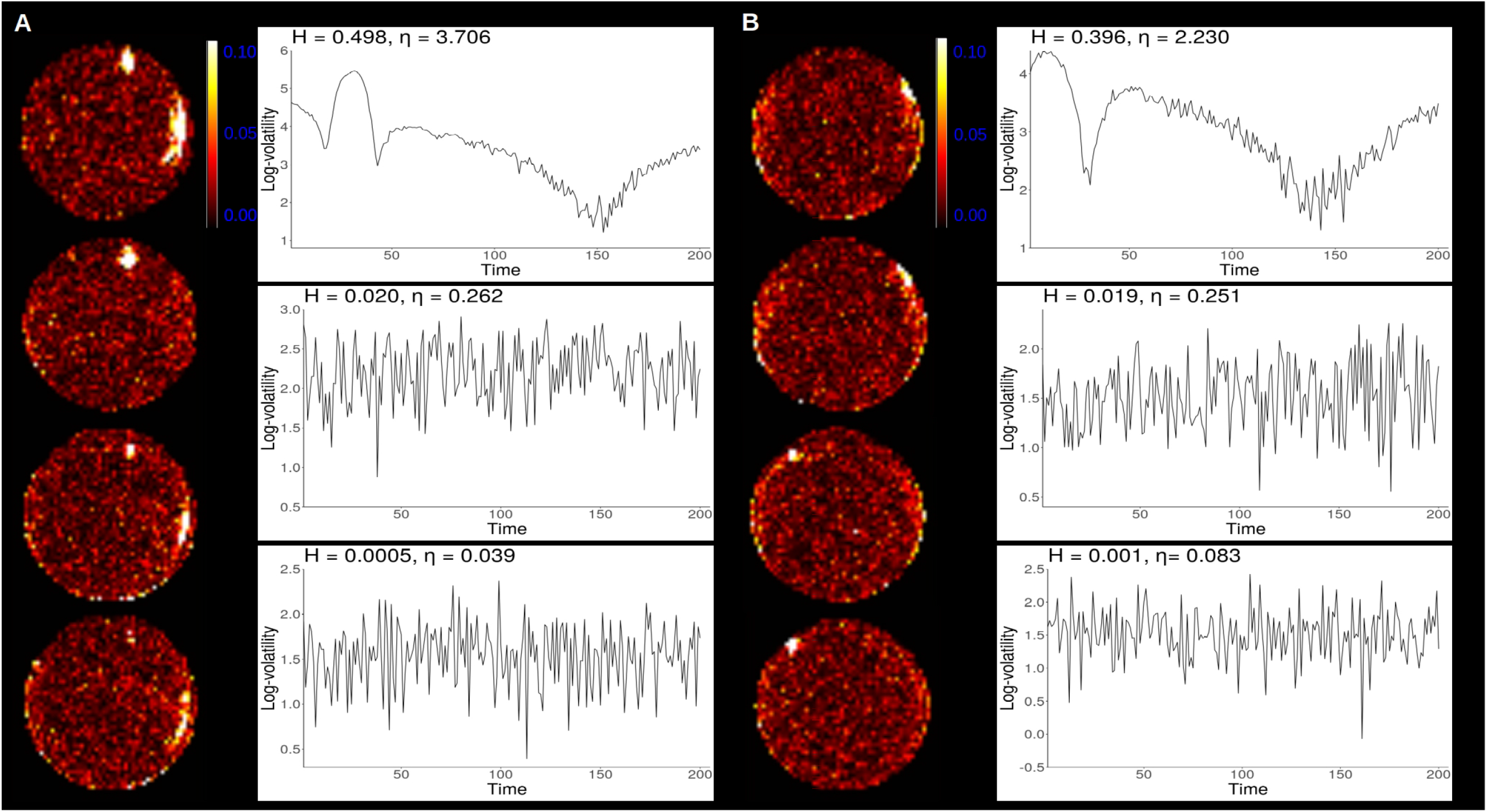
Spatial distribution of estimated *H* parameters in the phantom data Multi-slice view of scanner 1 phantom 1 (A) and scanner 2 phantom 2 (B) with log-volatility processes corresponding to maximum, minimum and mean *H* estimates.

**Figure 5:**
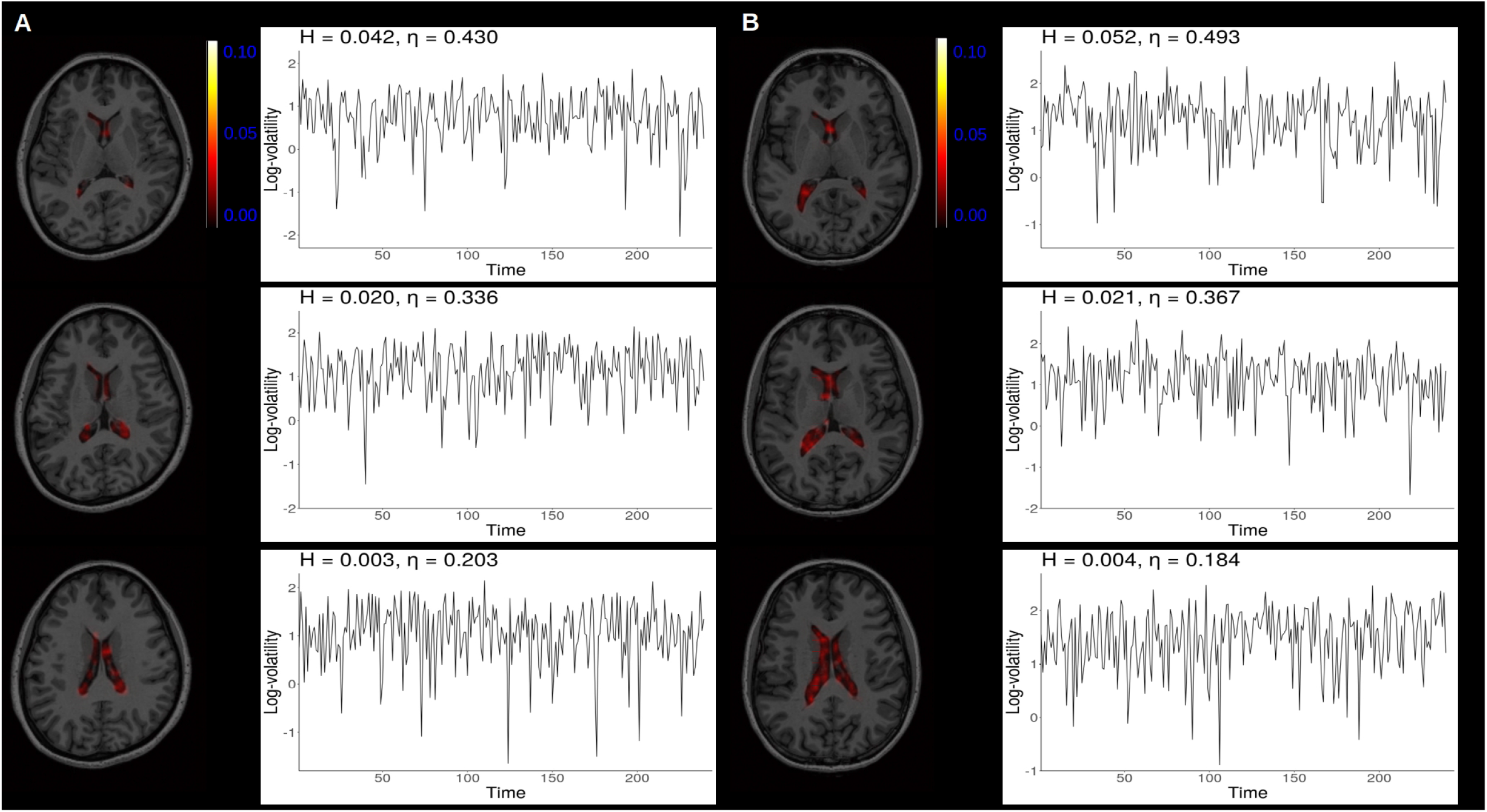
Spatial distribution of estimated *H* parameters in vivo noise Multi-slice view of data extracted from the ventricles of two participants, sub-17821 (A) and sub-21300 (B), from the ds000258 dataset with log-volatility processes corresponding to maximum, minimum and mean *H* estimates.

**Figure 6:**
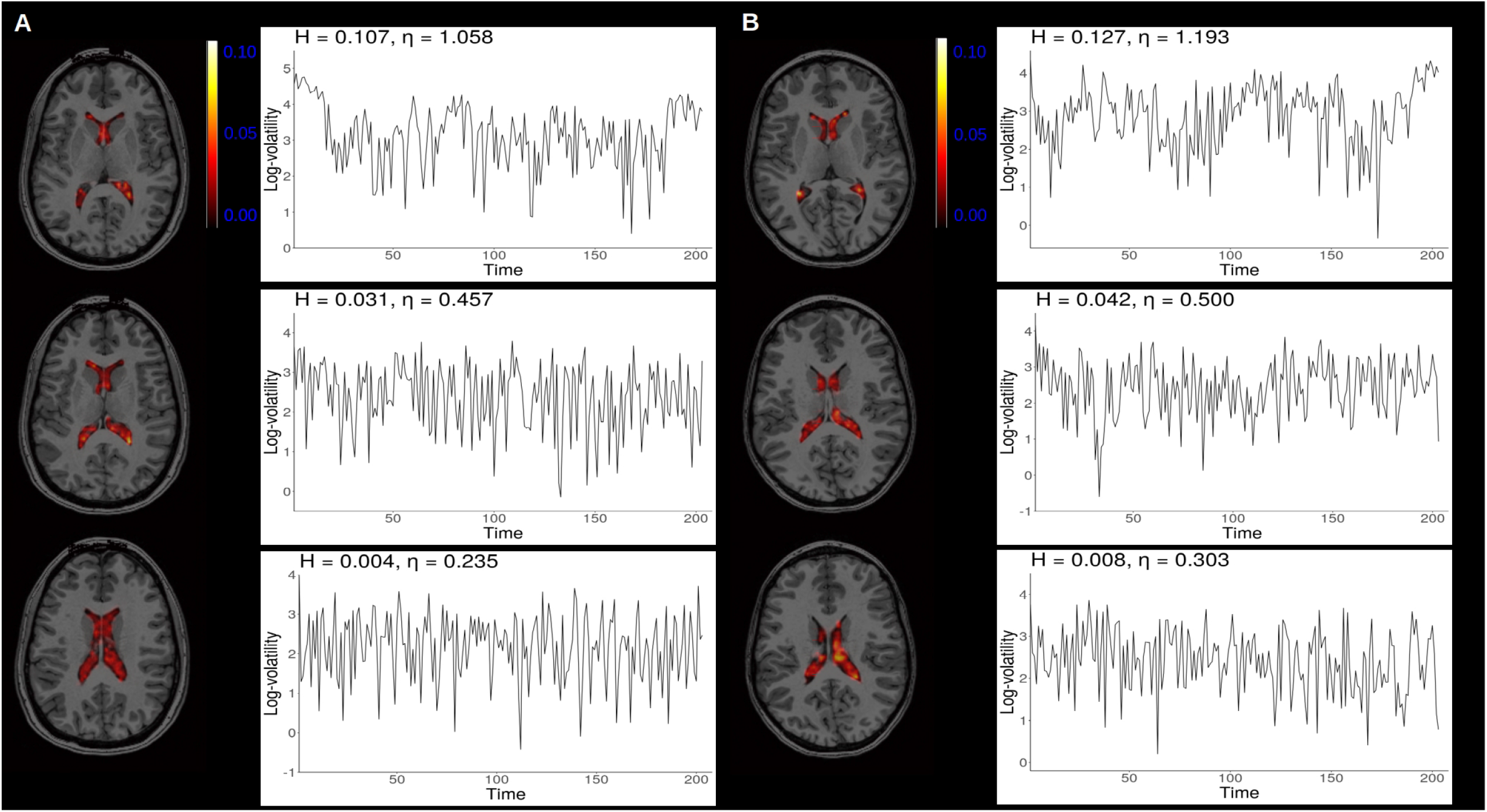
Spatial distribution of estimated *H* parameters in vivo noise Multi-slice view of data extracted from the ventricles of two participants, sub-28 (A) and sub-30 (B), from the ds000210 dataset with log-volatility processes corresponding to maximum, minimum and mean *H* estimates.

Some edge effects were observed, such that the estimated *H* parameters were generally larger near edges of the phantoms than the middle. In both phantoms the voxels with the maximum *H* parameter estimates were found near the edge and appeared to form large clusters. In the middle of the phantoms the *H* parameter estimates were generally very small yet varied. Similar edge effects were not observed in the resting state data extracted from the ventricles, which could be due to the fact that the ventricles reside close to the middle of the brain. Still, as in the phantom data, it was apparent that the *H* parameter estimates varied from voxel to voxel within the ventricles suggesting spatially non-constant volatility was present.

Interestingly, phantom 1 has a small region near the top where the signal intensity was lower than in the nearby voxels, suggesting signal dropout due to a possible air bubble (Supplementary Figure 14). This area consisted of four voxels and one of these voxels had the largest *H* parameter estimate in phantom 1. This voxel also represents the centre of the cluster near the top of the phantom in Figure 4A. No such signal dropout was seem in phantom 2.

### 3.3. Agreement between the correlation governed by the Hurst parameter H and ARFIMA autocorrelation

As mentioned earlier, the Hurst parameter not only governs the roughness of a volatility path, but also the autocorrelation function of the volatility time series. In this section we test for the agreement between the presence of autocorrelation predicted by the rough volatility model and the autocorrelation predicted by a standard ARFIMA model. As shown in Table 2 the correlation was significant and positive in both phantoms and data extracted from the ventricles.

**Table 2:** Correlations of *H* and *d* parameters from different sources.

|  | Phantom data |  | ds000528 |  | ds000210 |  |
| --- | --- | --- | --- | --- | --- | --- |
|  | Phantom 1 | Phantom 2 | sub-17821 | sub-21300 | sub-28 | sub-30 |
| $\rho$ | 0.69 | 0.69 | 0.40 | 0.58 | 0.68 | 0.79 |
| p-value | < 0.001 | 0.002 | < 0.001 | < 0.001 | < 0.001 | < 0.001 |

$\rho$ = Spearman correlation coefficient; ds000258 and ds000210 refer to the two Openneuro. datasets used.

The relationship between estimated *H* parameter and the ARFIMA[0, *d,* 0] memory parameter, *d*, of the log-volatility processes are presented in Supplementary Figures 7 and 8. The correlations between the *H* and *d* parameters was more variable in the resting state data extracted from the ventricles, which may be related to the fact that the size of the ventricles and thus the number of voxels in the the ventricles varied between participants and participants with more voxels inside the ventricles has higher correlations.

## 4. Discussion

The aim of the present empirical study was to examine the roughness of fMRI noise volatility. We used multi-echo scans of two phantoms from two different MRI scanners to estimate realised volatility. We also used human data from two separate multi-echo resting state datasets to examine whether patterns observed in the phantom data were present in vivo noise, specifically, focusing on signal extracted from the ventricles. Roughness of logarithm of the realised volatility processes was estimated using CNN calibration tools introduced in [31]. The findings indicated that no fMRI noise volatility series had constant volatility with estimated volatility of volatility *η* > 0. On average the volatility of fMRI noise is very rough with *H* ≈ 0.03, but substantial variability was also observed. The variability was caused by the fact that the smoothness of the volatility was not constant across the phantoms, with higher *H* estimates observed near the edges of the images. Interestingly, similar patterns of variability, with the exception of edge effects as we focused on data from the ventricles, were also observed in vivo noise, but the average was somewhat higher, *H* ≈ 0.05 and maximum substantially lower. Overall, all *H* < 0.5 suggesting that across the phantom and human data, volatility was consistently rough.

The present findings suggest that log-volatility of fMRI noise appears to behave like fractional Brownian motion with *H* parameter estimates between 0.03 and 0.05. As anticipated, these findings go some way to mimic the rough volatility pattern observed in high frequency financial data with Hurst parameter estimates varying between 0.02 and 0.14 [18, 17]. Thus, it appears that although fMRI scanner noise on average does not have large fluctuations in volatility over time, i.e. the noise does not exhibit sustained periods of high volatility followed by sustained periods of low volatility. Instead, the noise processes exhibit rapid spikes and oscillations, indicating more “severe” heteroscedasticity. The heteroscedasticity observed in the phantom data cannot be explained by head motion, physiology, or other known sources of non-constant noise and cannot be easily entered into analysis as a covariate because scanner noise processes cannot be directly observed during brain scanning. These findings challenge the assumption that fMRI noise has constant volatility and adds to the steady accumulation of literature exploring heteroscedasticity in fMRI noise [1, 2, 3, 4, 5, 6], further highlighting the importance of taking non-constant noise into consideration during analysis of the time series data.

The impact of rapidly spiking and oscillating volatility on fMRI data analysis has recently been investigated. One study examined the impact of heteroscedasticity introduced by simulated head motion spikes on fMRI data analysis [1]. The authors found that a linear modelling approach based on weighted least sum of squares (WLSS) was able to accurately model impulse responses to stimuli if the heteroscedasticity was constant across all voxels [1]. However, when the number of head motion spikes varied from voxel to voxel, the WLSS failed to accurately detect impulse responses. These findings led the authors to propose a heteroscedastic general linear model which incorporates head motion covariates. However, our findings suggest that not only can heteroscedasticity also be present in the scanner noise, but also the pattern of heteroscedasticity varies from voxel to voxel, with different patterns of spiking and rapid oscillations. Furthermore, our findings also indicate that similar patterns in volatility can be observed in the human data, which can be taken to suggest that the heteroscedasticity observed in scanner noise is also present in vivo noise. Taking the above findings by [1] into consideration, it is possible that such spatially non-constant heteroscedasticity in fMRI noise could influence data analysis.

Interestingly, to our knowledge only a few studies to date have examined the impact of heteroscedastic noise not explained by head motion or physiology on fMRI data analysis. In all studies the authors examined the usefulness of deterministic autoregressive conditional heteroscedasticity (ARCH) and generalised ARCH (GARCH) -type models, to aid investigation of time-dependent functional connectivity [6, 25, 41]. The studies specifically investigated GARCH(1,1) models with only one autoregressive and one moving average lag, suggesting the authors assumed the volatility would exhibit short memory. Simulation and real data experiments both showed that incorporating GARCH(1,1) model into the analysis helped to accurately model the time-dependent functional connectivity. Traditional approaches, including sliding window and exponentially weighted moving average models, on the other hand, were found to produce more false positive findings [6, 25, 41]. Moreover, previous Monte Carlo experiments have shown that heteroscedasticity violates the assumptions of not only correlation tests but also linear regressions in ways that can produce false positive findings [42, 43, 44]. Taken together with the present findings, we believe that further investigation of the impact of short memory heteroscedasticity on various different fMRI data analysis methods as well as selecting the most efficient and accurate methods to model the time-dependent volatility is of interest. Such further work could ultimately help improve both resting state and task-based data analysis as the noise in the time series is better understood [3, 45].

The present findings also show that the roughness of fMRI noise is not constant across regions in the phantom with the edges showing greater smoothness in the volatility relative to the centre of the phantom. This suggests that the volatility near the edges of the phantom was more likely to exhibit sustained periods of high and low volatility rather than rapid oscillations or spiking behaviour. To an extent these findings mirror those from previous work examining long-range dependence in the mean of fMRI noise [46, 47]. Previous studies have found that the long-range dependence near the edges of the phantom has estimated *H* > 0.5, indicating persistence and sustained periods of high and low mean in the series [46]. Similar edge effects have also been observed in real brain scans [48, 46]. Taken together with the present findings this suggests that fMRI data near the edges of an image appears to be more complex than that near the centre. Such time-dependent behaviour in the noise near the edges complicates data analysis as these effects violate assumptions of most time series modelling methods and can lead to both spurious regressions and correlations [49, 50, 51, 52, 53, 42, 43, 44]. Further investigation of the impact of reported edge effects on fMRI data analysis methods is of interest.

It is also important to note that in the present study, one of the phantoms had a small region of signal dropout, possibly indicating a presence of an air bubble. This region was the centre of one of the clusters where the smoothness of the volatility process was greater than in nearby regions. Previous studies have also found that air bubbles in phantoms can lead to drop in signal intensity, which has been suggested to due to susceptibility artifacts at the air-water boundary [54]. Air bubbles can also introduce phase errors and related magnetic field heterogeneity [55, 56], which could go some way to explain the larger *H* estimates in one of the clusters in one of the phantoms. Interestingly, such an effect was only found in one of the phantoms, suggesting that all the edge effects could not be explained by air bubbles. Still further investigation of the spatial pattern of volatility in fMRI noise in a gel phantom prepared with warm water, which are less susceptible to air bubbles [57], would be of interest.

The present study is not without limitations. First, the CNN was trained using simulated data as it was not possible use true realised volatility data in the training because the true *H* and *η* of such data are unknown. Although this method has been previously used in the field of financial mathematics and have been shown to outperform alternative models, such as those based on the sum of least squares [31], a model is always a simplification of reality. However, we argue that even though the simulated log-volatility paths used in the training of the CNN may indeed be different from the real data, they are no more different than the constant volatility assumption of traditional fMRI time series analysis methods. Additionally, we chose to use the mean reverting and driftless rBergomi model to simulate data because it closely reflects the behaviour in fMRI data. Additionally, the resting state data used to examine whether volatility patterns observed in the phantoms could also be seen in noise in vivo could have been influenced by head motion. Although, we took steps to minimise the impact of head motion on the analysis, it is possible that *H* parameters estimates were still influenced by head movements. However, considering the pattern of volatility observed in the resting state data extracted from the ventricles largely mirrored that seen in the middle of the phantoms we believe it can be concluded that at least some of the rapidly oscillating heteroscedastic scanner noise is present in vivo.

In the present study, realised volatility was estimated after slice timing correction and realignment, but no further preprocessing or de-noising steps were taken prior to estimation. This was done in an attempt to mirror standard multi-echo preprocessing pipelines where the echoes are normally combined prior to further preprocessing steps, such as smoothing, and independent component analysis-base de-noising [36, 34, 58]. This meant that we were unable to examine the impact of de-noising on realised volatility. Additionally, realised volatility was estimated using only eight echo time points as this was the maximum number we were able to collect. In finance, on the other hand, it is common to use high frequency asset price data, with sub-second granularity, to estimate daily volatility. It is difficult to ascertain whether our use of lower frequency data to estimate realised volatility had an impact on the present findings. Finally, the phantom and human data were acquired using different 3 Tesla MRI units. It is possible that the volatility of fMRI noise from scanners with different field strengths might vary and further investigation of this may be of interest.

## 5. Conclusions

The aim of the present study was to examine the smoothness of estimated realised volatility of fMRI noise as well as to examine whether patterns identified in the phantom scans were present in human data. This was done by conducting two multi-echo scans of two phantoms using two different MRI scanner units and using publicly available multi-echo resting state data. Multi-echo data were used to estimate realised volatility by *T*_2_∗-weighted variance. Smoothness of the realised volatility data was estimated by following cutting edge methods developed in the field of financial mathematics, namely by training a CNN to predict the Hurst parameter, *H*. The findings showed that on average scanner noise is very rough with *H* ≈ 0.03 and the roughness of the volatility data varied across the spatially across the phantoms. In both phantom scans the *H* estimates were larger near the edges, suggesting that volatility was smoother in these regions. Similar patterns of variability, with the exception of large edge effects, were observed in the resting state data extracted from the ventricles. Thus, seems that rapidly oscillating, spatially non-constant heteroscedastic noise is present in vivo noise as well. Taken together the present findings further challenge the assumption that fMRI scanner noise has constant volatility and highlight the need for further research to investigate how to effectively model the heteroscedasticity during time series analysis.

## 6. Acknowledgements

JL is supported by Sir Henry Wellcome Postdoctoral Fellowship (213578/Z/18/Z). The funding body did not play an active role in the design of this study, nor in data collection or analysis, nor in writing the manuscript.

## Appendix A. Artificial Neural Networks

In this appendix we repeat the introduction given by Stone [31, Section 2.1, p382] for the reader’s convenience. An artificial neural network is a biologically inspired system of interconnected processing units, where each processing unit is called a layer. Inputs to each layer, apart from the first layer, are outputs from previous layers. A layer is composed of a number of nodes, and each node in a given layer is connected to the nodes in a subsequent layer, thus forming a network; each edge in this network has a weight associated to it. The first processing unit is called the input layer, and the final processing unit is the output layer. The processing unit or units between the input layer and output layer are referred to as hidden layers; typically artificial neural networks have more than one hidden layer.

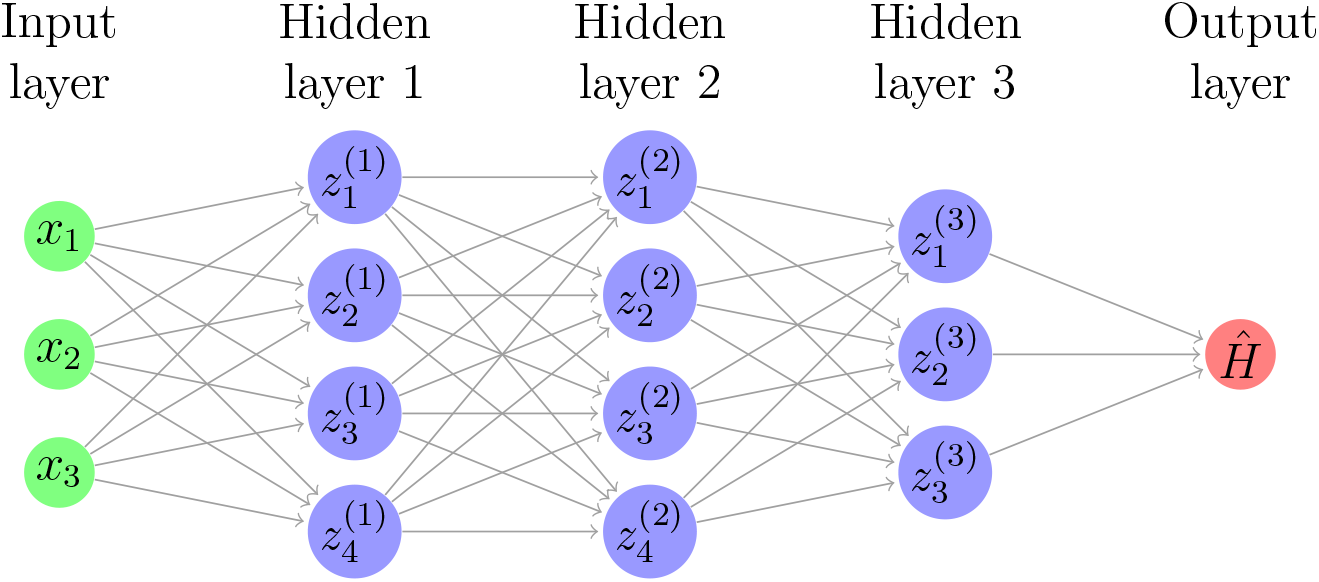

Convolutional neural networks (CNNs) are a class of artificial neural networks, where the hidden layers can be grouped into different classes according to their purpose; one such class of hidden layer is the eponymous convolutional layer. Below we describe the classes of hidden layers used in our CNN. Of course, this list is not exhaustive, and there exist many classes of hidden layers that we omit for means of brevity. Note also that we describe a CNN in the context of the problem we are trying to solve, where the input data are one dimensional vectors. CNNs can of course also be used on higher dimensional input data, but the fundamental structure and different roles of the hidden layers do not change.

- **Convolutional Layer:** In deep learning, the convolution operation is a method used to assign relative value to entries of input data, in our case one dimensional vectors of time series data, while simultaneously preserving spatial relationships between individual entries of input data. For a given kernel size *k* and an input vector of length *m*, the convolution operation takes entries 1,…, *k* of the input vector and multiplies by the kernel element-wise, whose length is *k*. The sum of the entries of the resulting vector are then the first entry of the feature map. This operation is iterated *m* + 1 − *k* times, thus incorporating every entry in the input data vector into the convolution operation. The output of the convolutional layer is the feature map. For example, let (1, 2, 1, 0, 0, 3) be our input vector, and (1, 0, 1) be our kernel; here the kernel size is 3. The first iteration of the convolution operation involves taking the element-wise multiple of (1, 2, 1) and (1, 0, 1): (1, 0, 1) is produced and the sum, equal to 2, is computed. This is the first entry of the feature map. The resulting feature map in this example is then (2, 2, 1, 3). Clearly, the centre of each kernel cannot overlap with the first and final entry of the input vector. Zero-padding, sometimes referred to as same-padding, preserves the dimensions of input vectors and allows more layers to be applied in the CNN: zero-padding is simply the extension of the input vector and the setting of the first and final entries as 0, while leaving the other entries unchanged. In our example, the input vector becomes (0, 1, 2, 1, 0, 0, 3, 0) after zero padding.
- **Activation Layer:** The activation layer is a non-linear function *σ* that is applied to the output of the convolutional layer i.e. the feature map; the purpose of the activation layer is indeed to introduce non-linearity into the CNN. Examples of activation functions include the sigmoid function and the hyperbolic tangent function. In our CNN we use the ‘LeakyReLU’ activation function, defined as

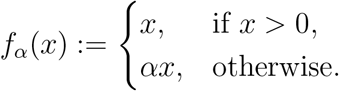

The LeakyReLU activation function allows a small positive gradient when the unit is inactive.
- **Max Pooling Layer:** For a given pooling size *p*, the max pooling layer returns a vector whose entries are the maximum among the neighbouring *p* entries in the feature map. For example, for feature map (1, 3, 8, 2, 1, 0, 0, 4, 6, 1) and *p* = 3 the max pooling output is (8, 8, 8, 8, 8, 2, 4, 6, 6, 6). Other pooling techniques apply the same idea, but use different functions to evaluate the neighbouring *p* entries in the feature map. Examples include average pooling, and L2-norm pooling, which in fact uses the Euclidean norm in mathematical nomenclature.
- **Dropout Layer:** Dropout is a well-known technique incorporated into CNNs in order to prevent overfitting. Without the addition of a dropout layer, each node in a given layer is connected to each node in the subsequent layer; dropout temporarily removes nodes from different layers in the network. The removal of nodes is random and determined by the dropout rate *d*, which gives the proportion of nodes to be temporarily dropped. Note that dropout is only implemented during training; during testing the weights of each node are multiplied by the dropout rate *d*.
- **Dense Layer:** Also referred to as the fully connected layer, each node in the input layer is connected to each node in the output layer as the name suggests. After being processed by the convolutional, activation, pooling, and dropout layers, the extracted features are then mapped to the final outputs via a subset of the dense layer, an activation function is then applied subsequently. This activation function is chosen specifically for the task that the CNN is required to execute, i.e. binary/multi-class classification, or regression to output a continuous value. The final output from the dense layer has the same number of nodes as the number of classes in the output data.

## Footnotes

1 This Brownian assumption implies a Hurst parameter of 0.5 (which is a parameter governing the Hölder regularity of paths) of the stochastic process. It is known, that for standard Brownian motion *H* = 0.5.

