## Supplementary Materials for "Sailing in rough waters: examining volatility of fMRI noise"

September 25, 2020

#### 1 Decay in fMRI noise

Figures 1, 2, 3, 4, 5, and 6 shows the pattern of decay from the first echo to the last at each time point in the time series extracted from five voxels from the phantoms and inside the ventricles from sub-17821 and sub-21300 from ds000258 as well as from sub-28 and sub-30 from ds000210.

Figure 1: Rate of decay from the first echo to the last in phantom 1

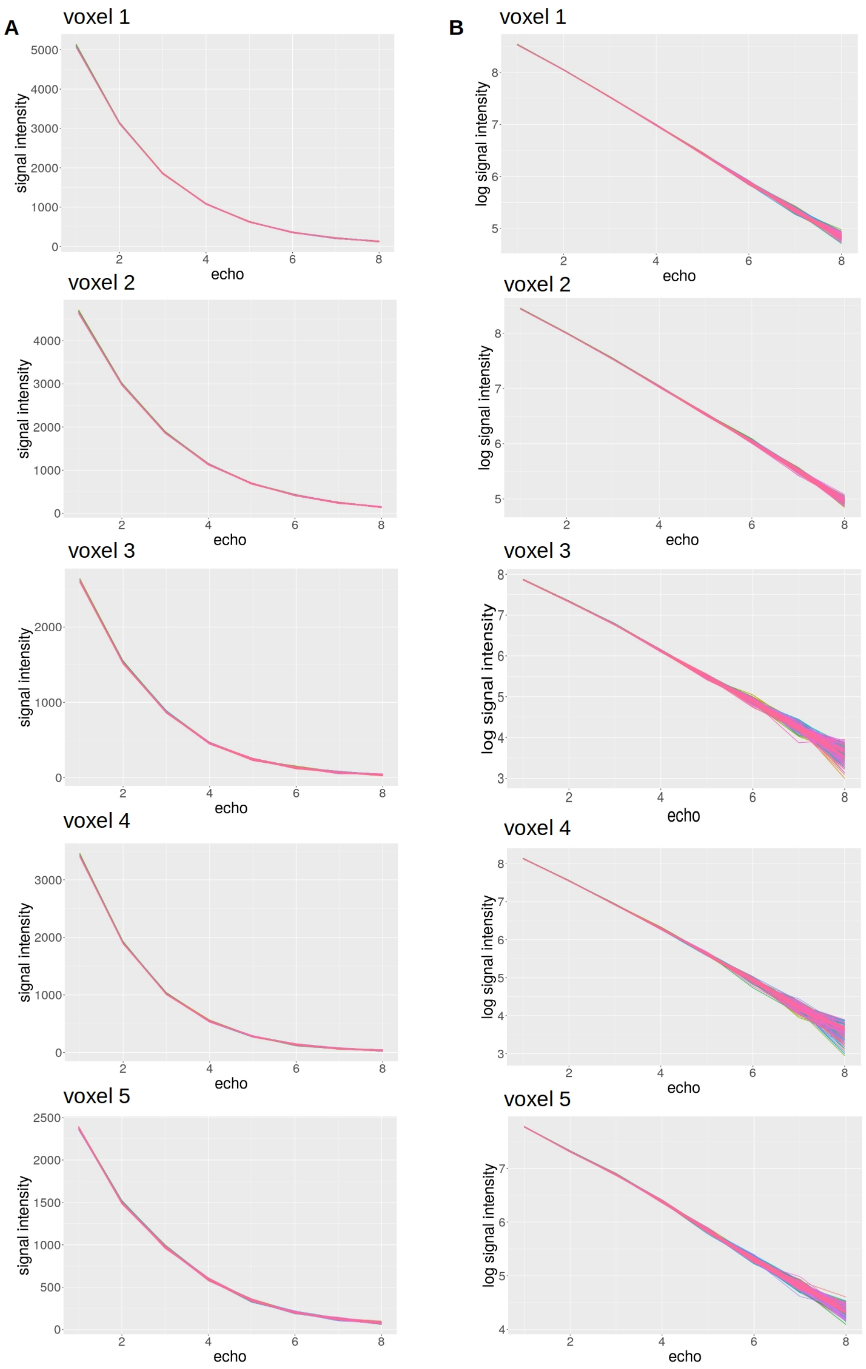


A. Raw signal intensity at each echo time; B. Logarithm transformed signal intensity at each echo.

Figure 2: Rate of decay from the first echo to the last in phantom 2

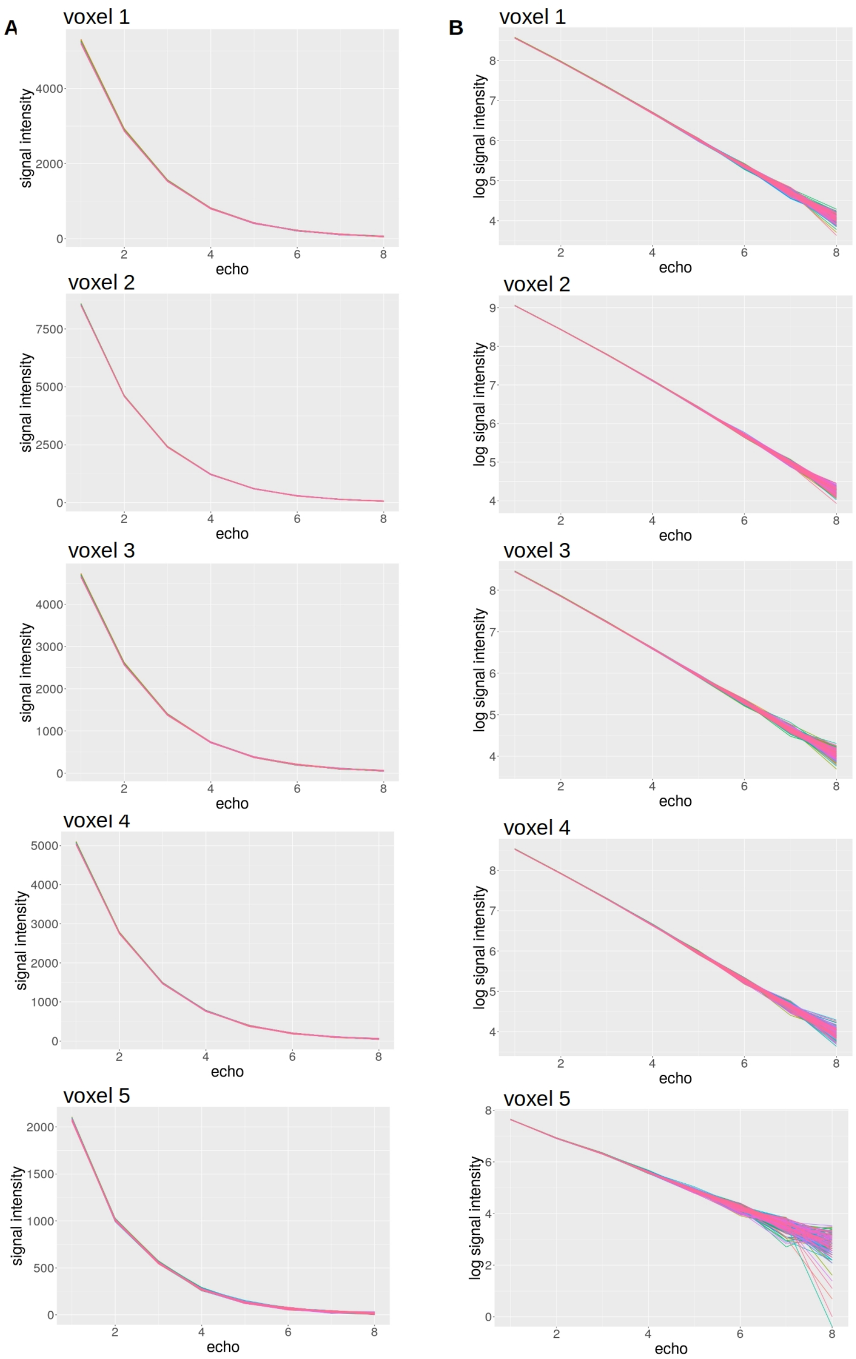


A. Raw signal intensity at each echo time; B. Logarithm transformed signal intensity at each echo.

Figure 3: Rate of decay from the first echo to the last sub-17821

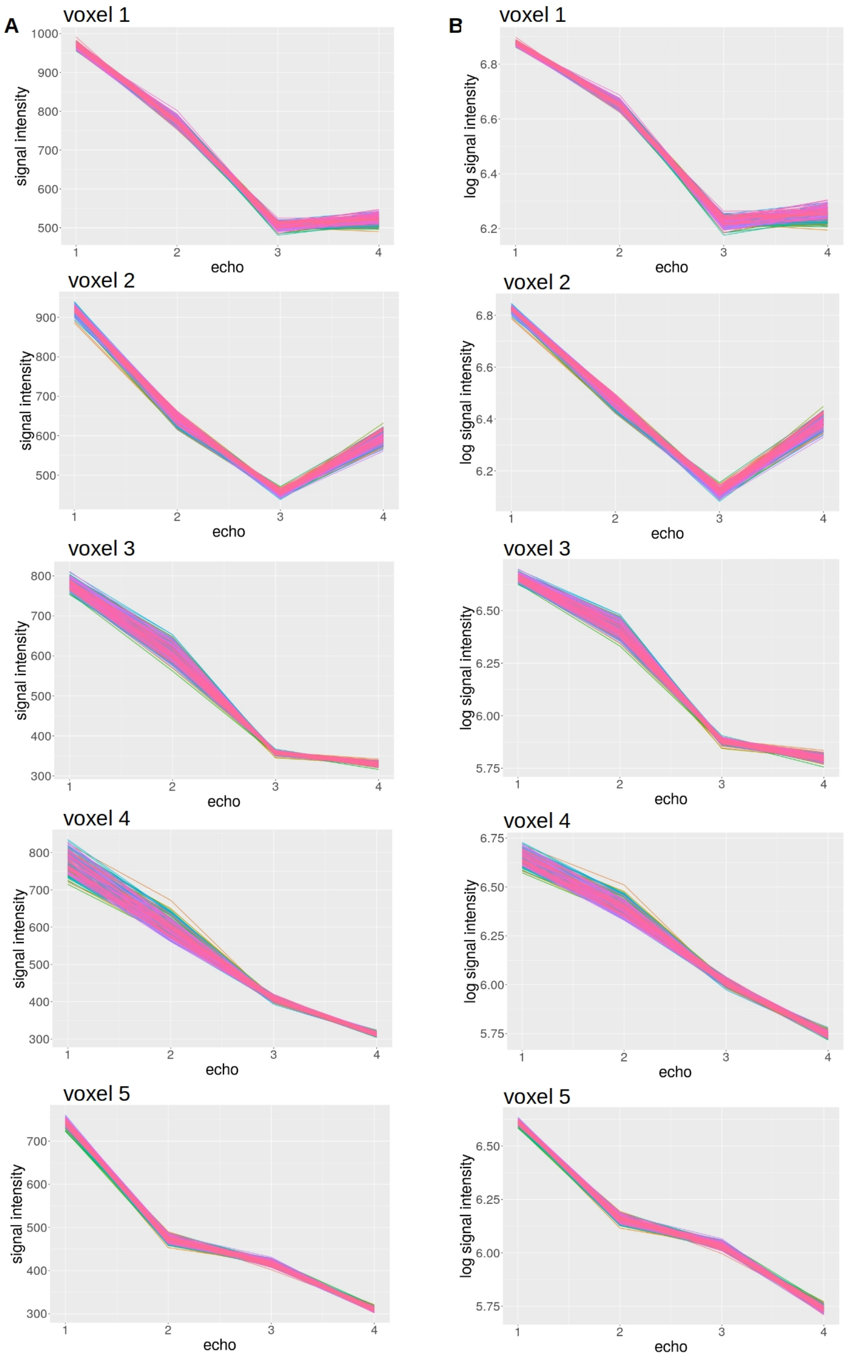


A. Raw signal intensity at each echo time; B. Logarithm transformed signal intensity at each echo.

Figure 4: Rate of decay from the first echo to the last sub-21300

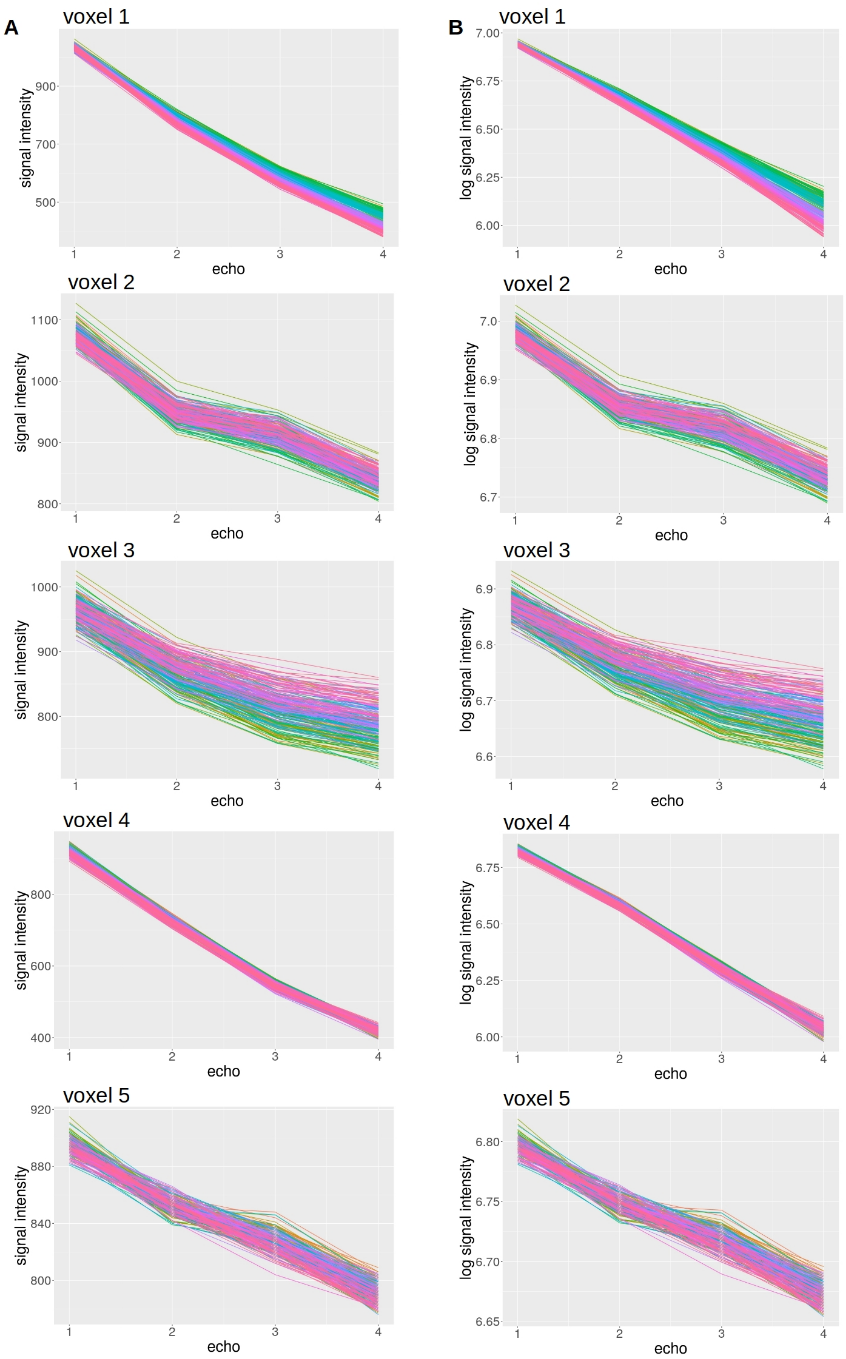


A. Raw signal intensity at each echo time; B. Logarithm transformed signal intensity at each echo.

Figure 5: Rate of decay from the first echo to the last sub-28

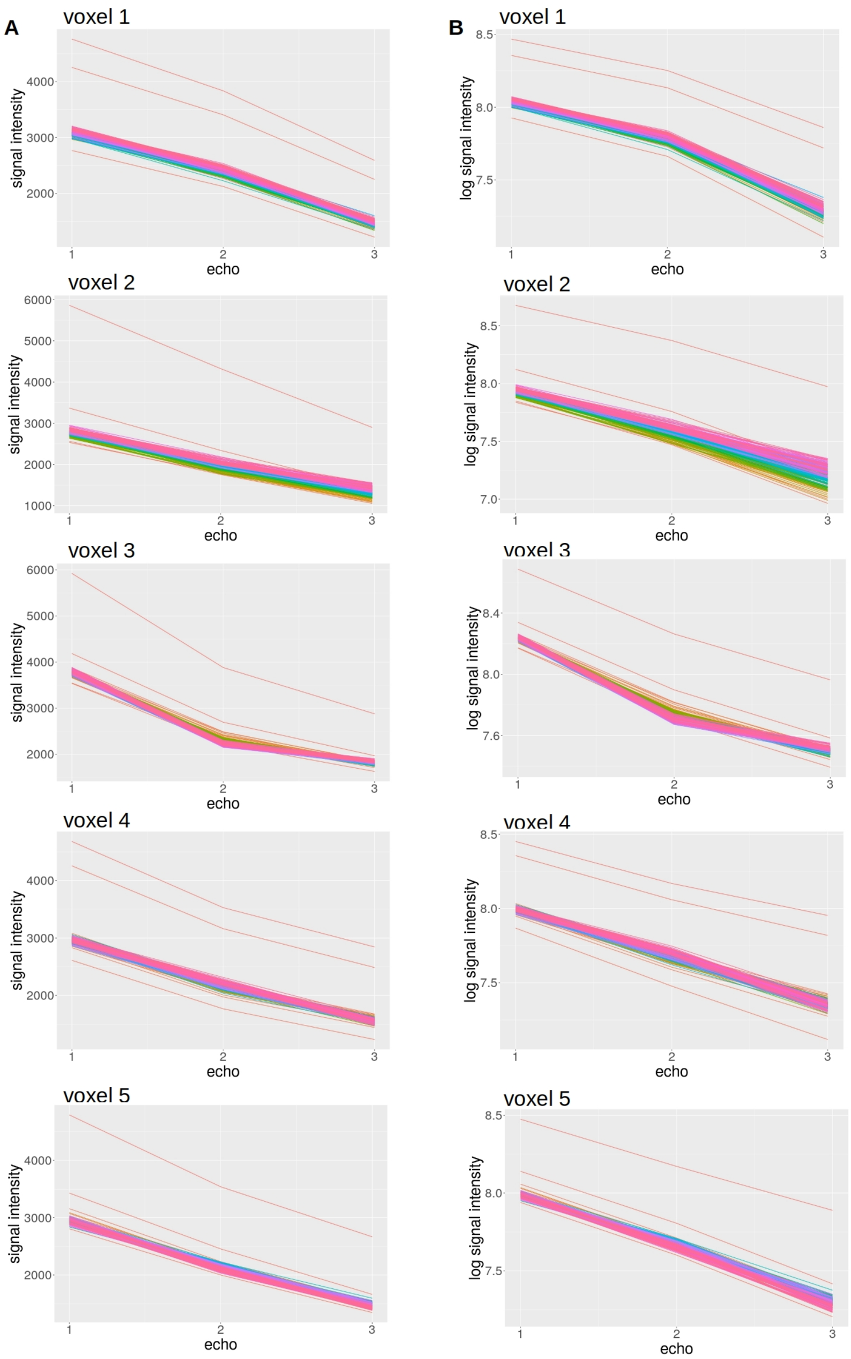


A. Raw signal intensity at each echo time; B. Logarithm transformed signal intensity at each echo.

Figure 6: Rate of decay from the first echo to the last sub-30

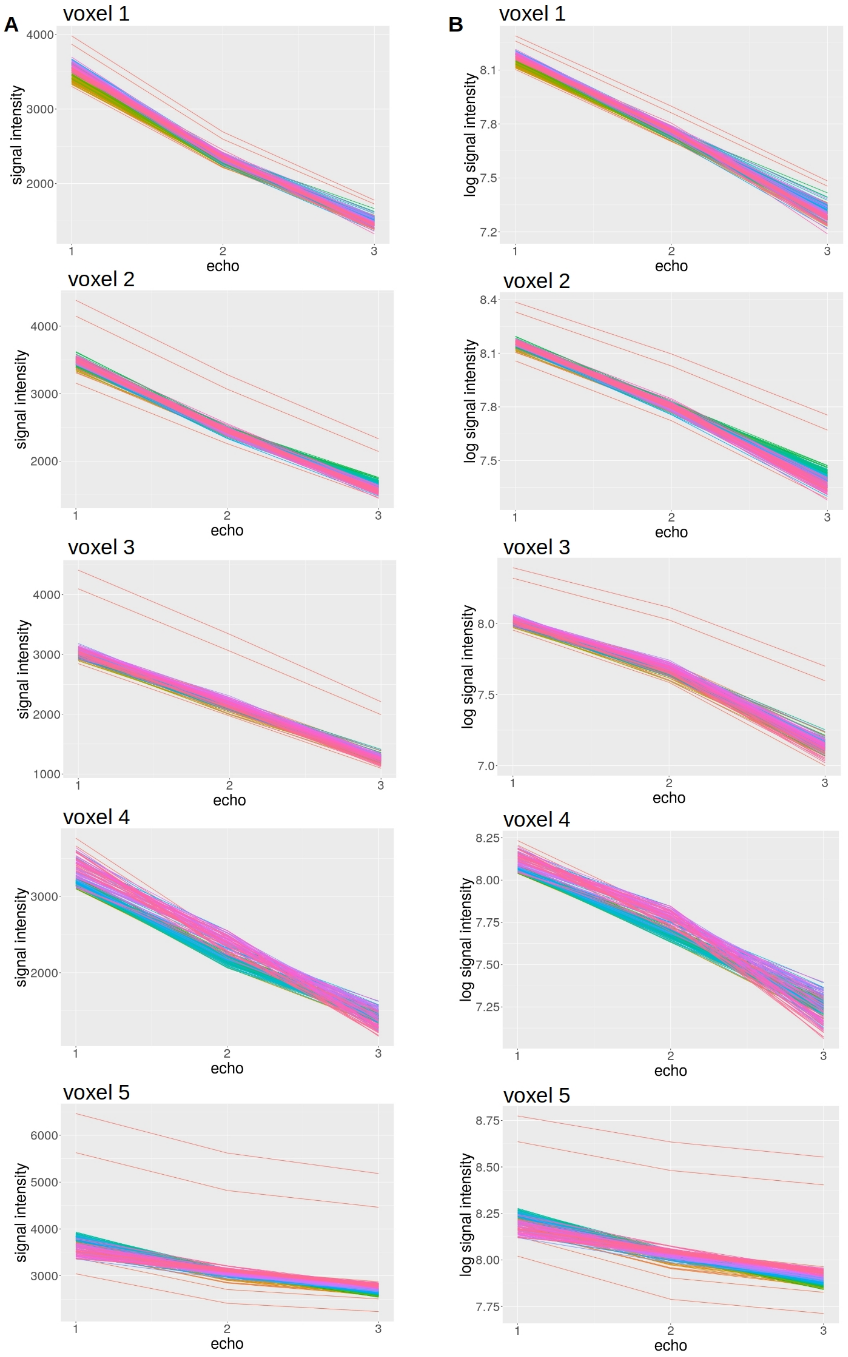


A. Raw signal intensity at each echo time; B. Logarithm transformed signal intensity at each echo.

### 2 Framewise displacement

Table 1: Framewise displacement summary statistics

| | | sub-17821 | sub-21300 | sub-28 | sub-30 |
|---|---|---|---|---|---|
| echo 1 | Mean | 0.049 | 0.052 | 0.049 | 0.063 |
| | SD | 0.027 | 0.034 | 0.043 | 0.049 |
| | Max | 0.228 | 0.294 | 0.464 | 0.523 |
| echo 2 | Mean | 0.061 | 0.063 | 0.059 | 0.081 |
| | SD | 0.031 | 0.036 | 0.039 | 0.054 |
| | Max | 0.239 | 0.278 | 0.450 | 0.543 |
| echo 3 | Mean | 0.063 | 0.062 | 0.061 | 0.093 |
| | SD | 0.030 | 0.037 | 0.037 | 0.058 |
| | Max | 0.229 | 0.294 | 0.450 | 0.481 |
| echo 4 | Mean | 0.063 | 0.065 | NA | NA |
| | SD | 0.030 | 0.036 | NA | NA |
| | Max | 0.227 | 0.310 | NA | NA |

Data from sub-17821 and sub-21300 came from dataset ds000258. Data from sub-28 and sub-30 came from dataset ds000210 and only 3 echo times were used. SD = standard deviation.

### 3 Relationship between the correlation governed by the Hurst parameter $H$ and ARFIMA autocorrelation

As can be seen in the figures, the log-volatility processes with larger estimated $H$ had substantially longer memory than log-volatility processed with very small estimated $H$. Some log-volatility series with small $H$ estimates still showed long memory, $d > 0$, but some were anti-persistent with $d < 0$. This suggests that the performance of the model was acceptable, providing accurate $H$ estimates.

Figure 7: Relationship between ARFIMA memory parameters $d$ and estimated $H$ parameter in phantom data

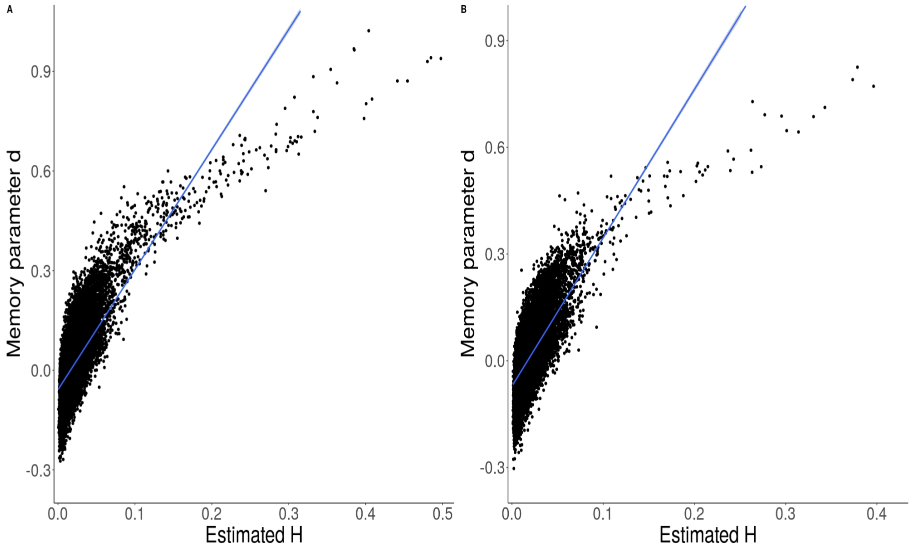


A. Phantom 1; B. Phantom 2. $d$ = ARFIMA$[0, d, 0]$ memory parameter, $H$ = Hurst parameter

Figure 8: Relationship between ARFIMA memory parameters $d$ and estimated $H$ parameter in vivo noise

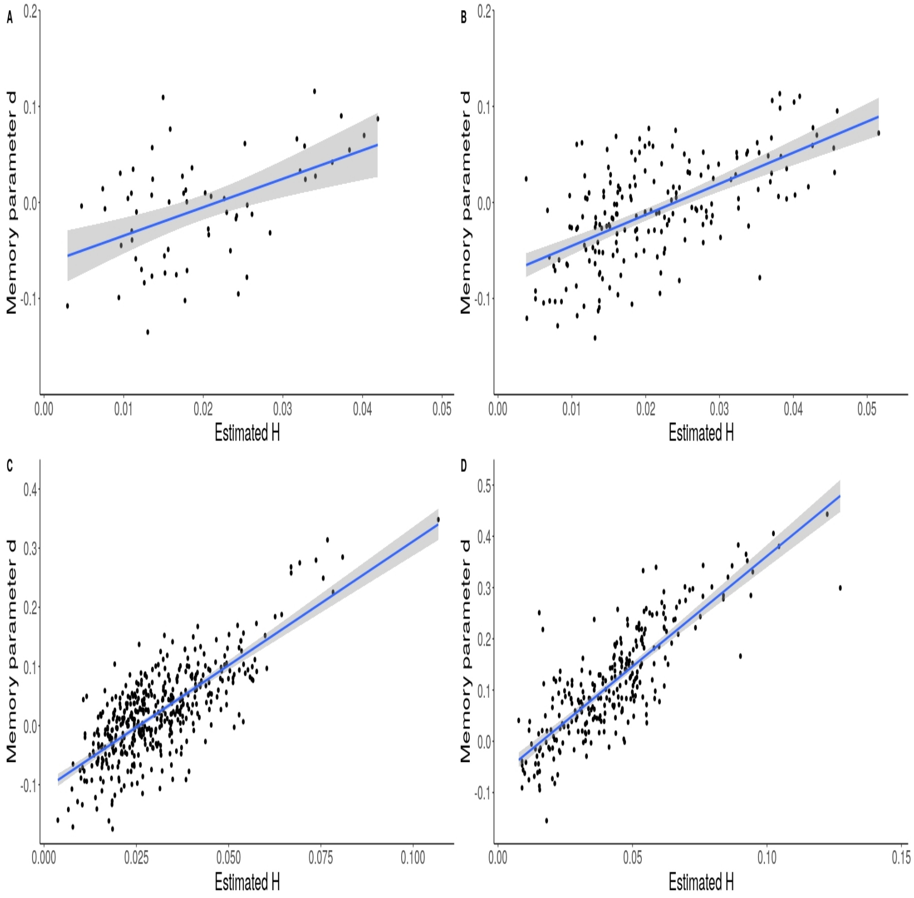


A. sub-17821 from ds000258; B. sub-21300 from ds000258, C. sub-28 from ds000210, D. sub-30 from ds000210. The number of datapoints is determined by the number of voxels inside the ventricles, which is determined by the size of the participant's ventricles. $d$ = ARFIMA$[0, d, 0]$ memory parameter, $H$ = Hurst parameter

### 4 Estimating roughness of fMRI noise using non-weighted realised volatility

#### 4.1 Estimating non-weighted realised volatility

The present study focused on studying volatility $v_t$ in fMRI noise. Multi-echo phantom data was used to study scanner noise and multi-echo resting state data was extracted from the ventricles of four participants to study combination of scanner and physiological noise. Thus, the data used in the present study was assumed to not contain any true brain signal, it was defined as:

$$y_t = \sqrt{v_t}\epsilon_t \tag{1}$$

where $t$ is the time point in the series and $\epsilon_t$ is the random white noise process.

Observations $y_t$ from each echo time $n$ up to the last echo $N$ were treated as intra-TR data which were used to estimate realised volatility for each point in the time series. The echos were combined and realised volatility were estimated by calculating sample mean and variance as follows:

$$\bar{x}_t = \frac{\sum_{n=1}^{N} y_{n,t}}{N} \tag{2}$$

Realised volatility at each time point,$t = 1...T$, was estimated by calculating variance between observations at each echo time $n$.

$$\hat{v_t} = \frac{\sum_{n=1}^{N} (y_{n,t} - \bar{x}_t)^2}{N} \tag{3}$$

The realised volatility paths were used to estimate the Hurst parameters $H$ using the same convolutional neural network tool described in the main manuscript.

#### 4.2 Estimated roughness and volatility of volatility

The summary statistics of the estimated $H$ and $\eta$ parameters of the realised log-volatility series in phantom and human data from the ventricles are presented in Table 2.

#### 4.3 Spatial pattern in estimated Hurst parameters

Figure 9 shows how the estimated $H$ varied from region to region across the phantoms and Figures 10 and 11 show the pattern of estimate $H$ parameter estimates in the ventricles. Log-volatility series associate with the maximum, minimum, and $H$ parameter estimates close to the mean are also shown.

Figure 9: Spatial distribution of estimated $H$ parameters in the phantom data

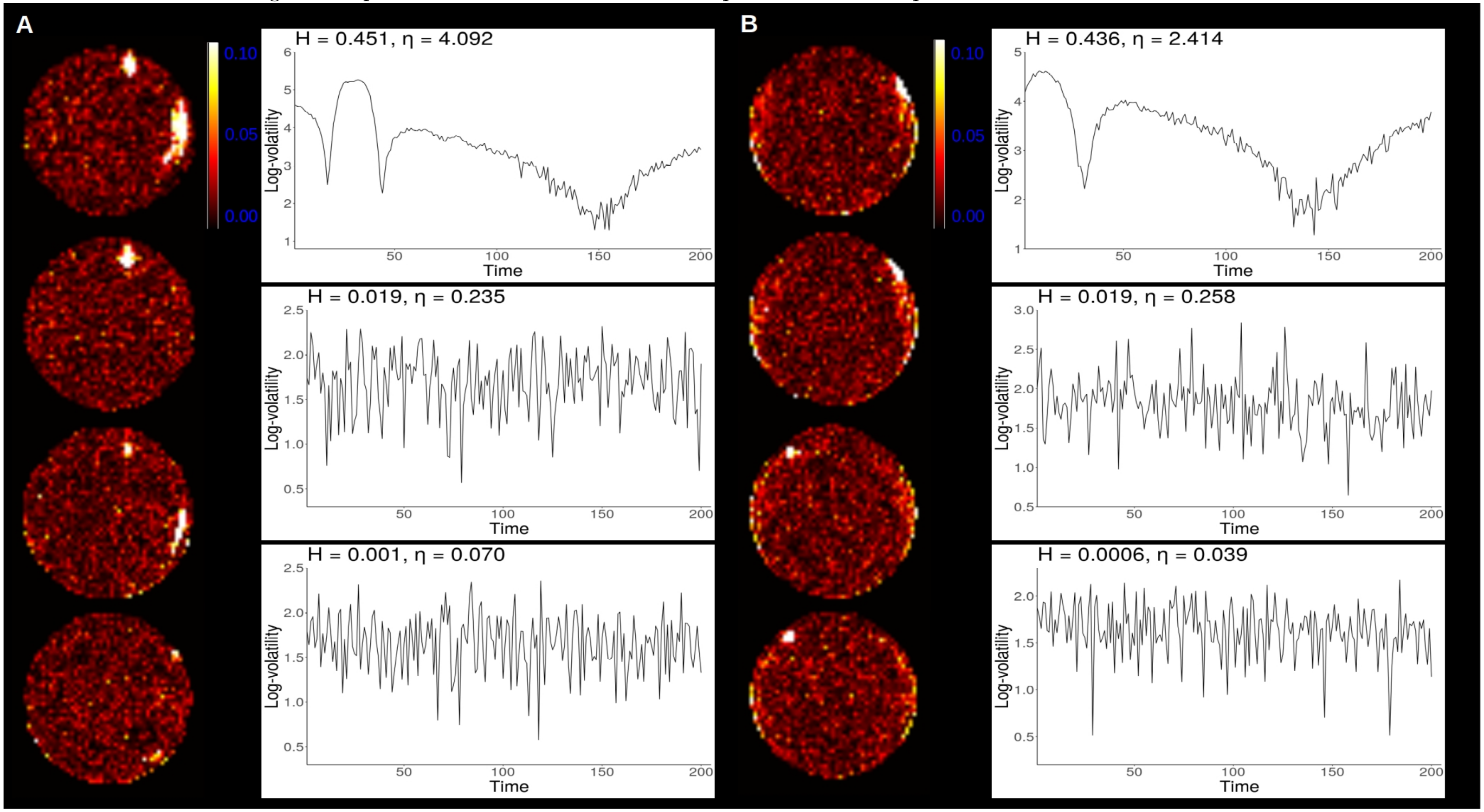


Multi-slice view of scanner 1 phantom 1 (A) and scanner 2 phantom 2 (B) with log-volatility processes corresponding to maximum, minimum and mean $H$ estimates.

Figure 10: Spatial distribution of estimated $H$ parameters in vivo noise

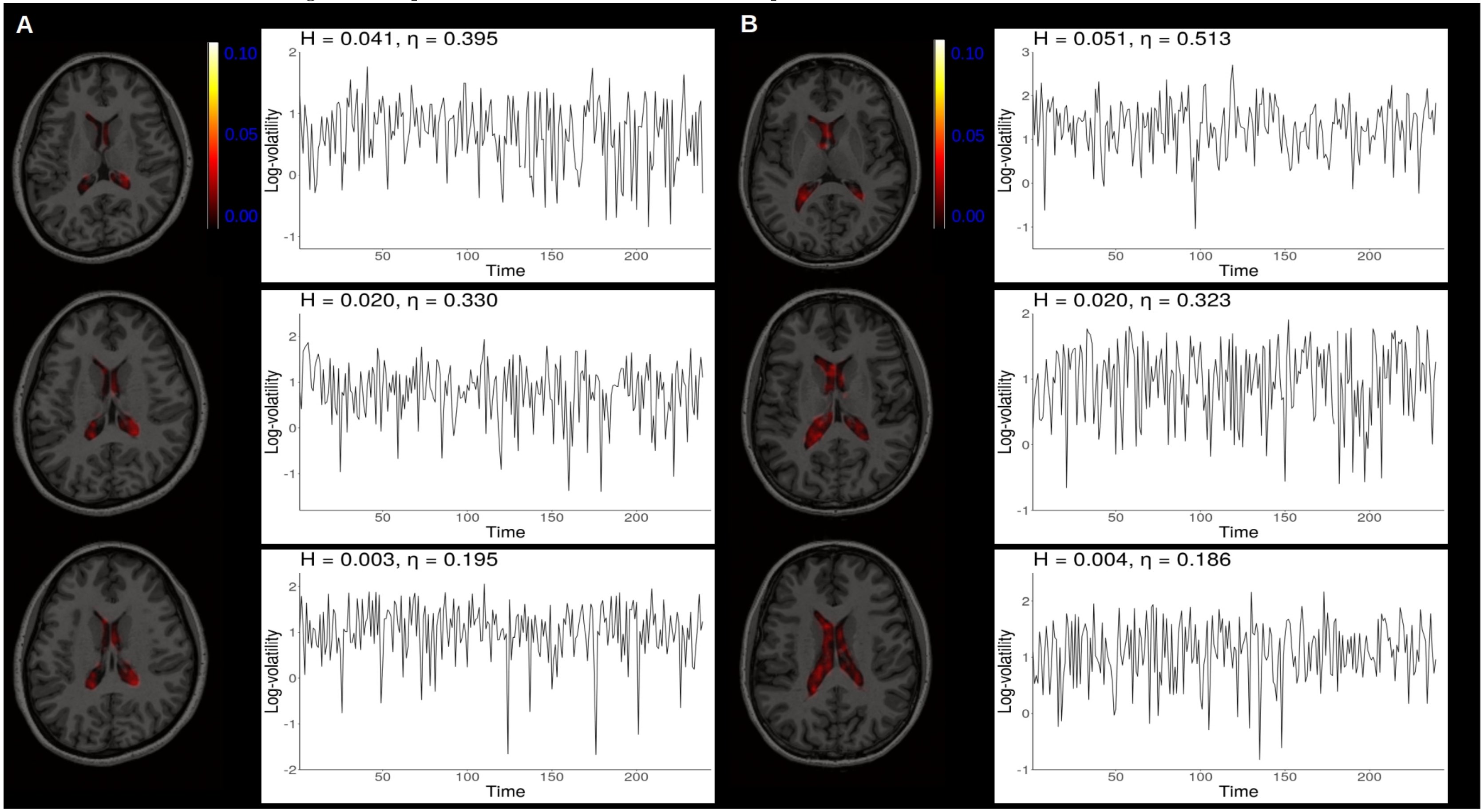


Multi-slice view of data extracted from the ventricles of two participants, sub-17821 (A) and sub-21300 (B), from the ds000258 dataset with log-volatility processes corresponding to maximum, minimum and mean $H$ estimates.

Figure 11: Spatial distribution of estimated $H$ parameters in vivo noise

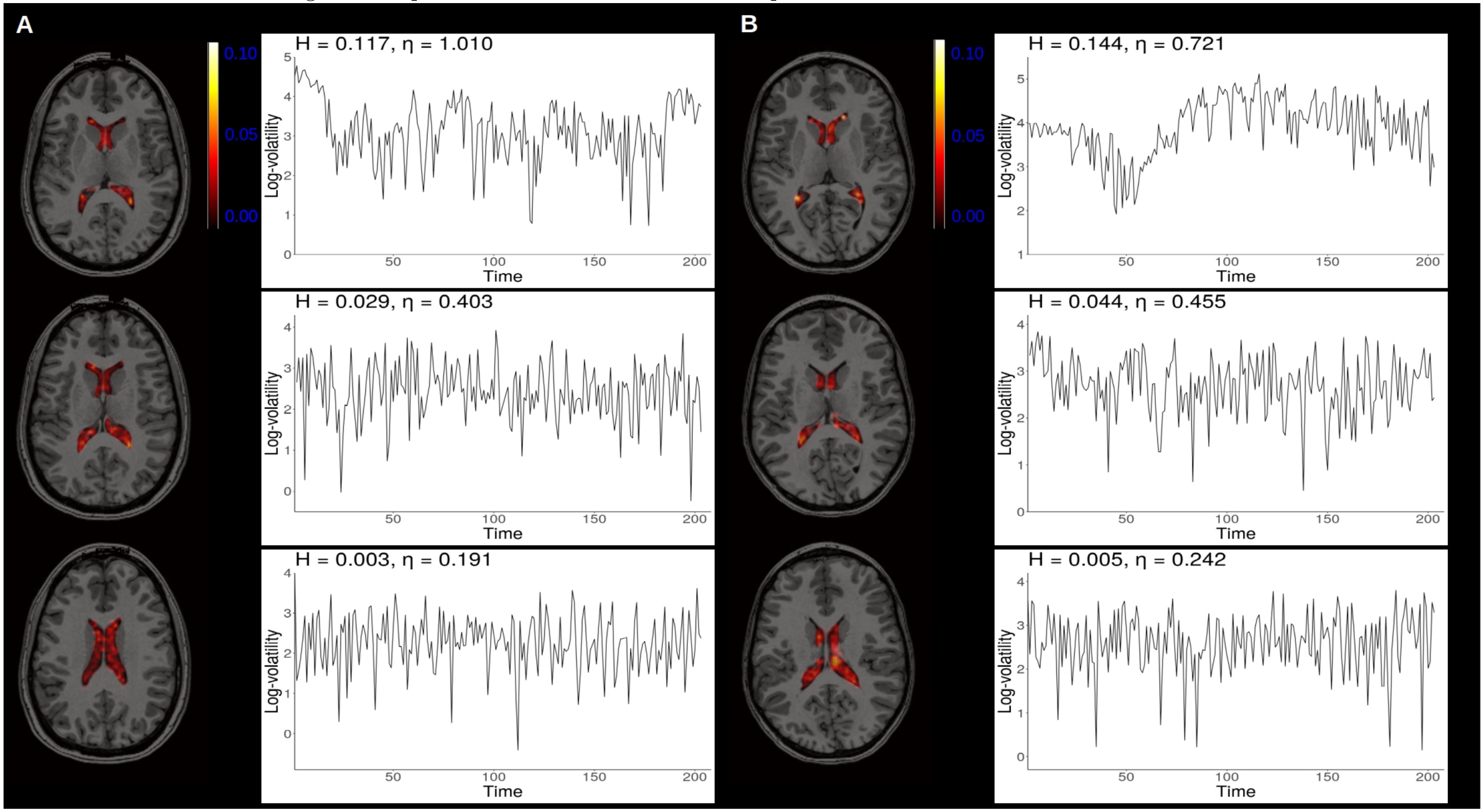


Multi-slice view of data extracted from the ventricles of two participants, sub-28 (A) and sub-30 (B), from the ds000210 dataset with log-volatility processes corresponding to maximum, minimum and mean $H$ estimates.

Table 2: Estimated $H$ and $\eta$ parameters in phantom and human data

| | | Phantom data | | ds000528 | | ds000210 | |
|---|---|---|---|---|---|---|---|
| | | Phantom 1 | Phantom 2 | sub-17821 | sub-21300 | sub-28 | sub-30 |
| $H$ | Mean | 0.019 | 0.019 | 0.020 | 0.019 | 0.044 | 0.029 |
| | SD | 0.020 | 0.017 | 0.010 | 0.009 | 0.023 | 0.015 |
| | Max | 0.451 | 0.436 | 0.051 | 0.041 | 0.144 | 0.117 |
| | Min | 0.001 | 0.0006 | 0.004 | 0.003 | 0.005 | 0.003 |
| $\eta$ | Mean | 0.224 | 0.211 | 0.322 | 0.309 | 0.479 | 0.410 |
| | SD | 0.113 | 0.077 | 0.054 | 0.049 | 0.129 | 0.087 |
| | Max | 4.092 | 3.950 | 0.513 | 0.400 | 1.186 | 1.010 |
| | Min | 0.039 | 0.039 | 0.186 | 0.195 | 0.233 | 0.191 |

Human data was extracted from the ventricles. SD = standard deviation; ds000258 and ds000210 refer to the two Openneuro. datasets used.

##### 4.4 Agreement between the correlation governed by the Hurst parameter $H$ and autocorrelation

The relationship between estimated $H$ parameter and the ARFIMA$[0, d, 0]$ memory parameter, $d$, of the log-volatility processes are presented in Figures 12 and 13.

The correlation was significant and positive in both phantoms: $r = 0.42, p < 0.001$ and $r = 0.56, p < 0.001$ for phantoms 1 and 2, respectively. Similarly, all correlations suggested a statistically significant positive relationship in ventricle data: $r = 0.043, p < 0.001$, $r = 0.58, p < 0.001$, $r = 0.67, p < 0.001$, and $r = 0.76, p < 0.001$ for sub-17821, sub-21300, sub-28, and sub-30, respectively.

##### 4.5 Air bubble in phantom data

Figure 14 shows a possible air bubble in phantom 1. No such air bubble was seen in phantom 2.

Figure 12: Relationship between ARFIMA memory parameters $d$ and estimated $H$ parameter in phantom data

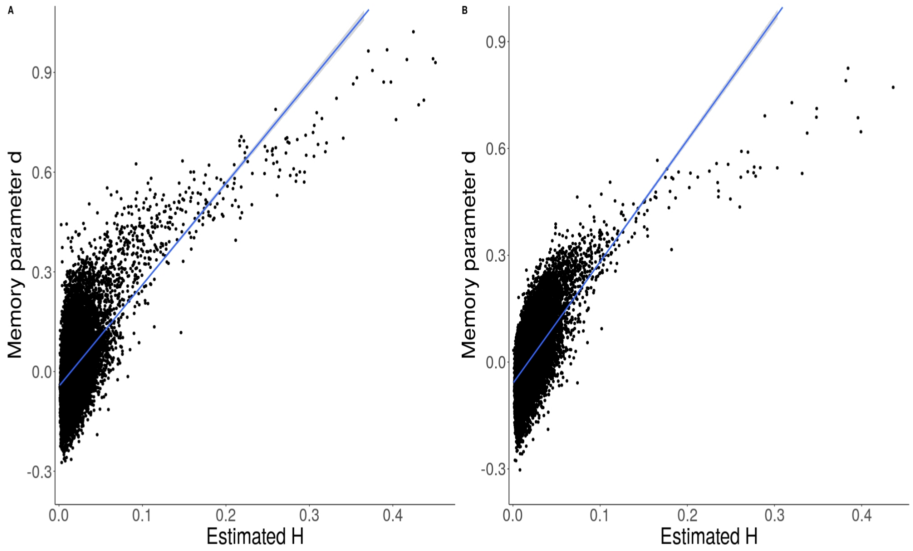


A. Phantom 1; B. Phantom 2. $d$ = ARFIMA$[0, d, 0]$ memory parameter, $H$ = Hurst parameter

Figure 13: Relationship between ARFIMA memory parameters $d$ and estimated $H$ parameter in vivo noise

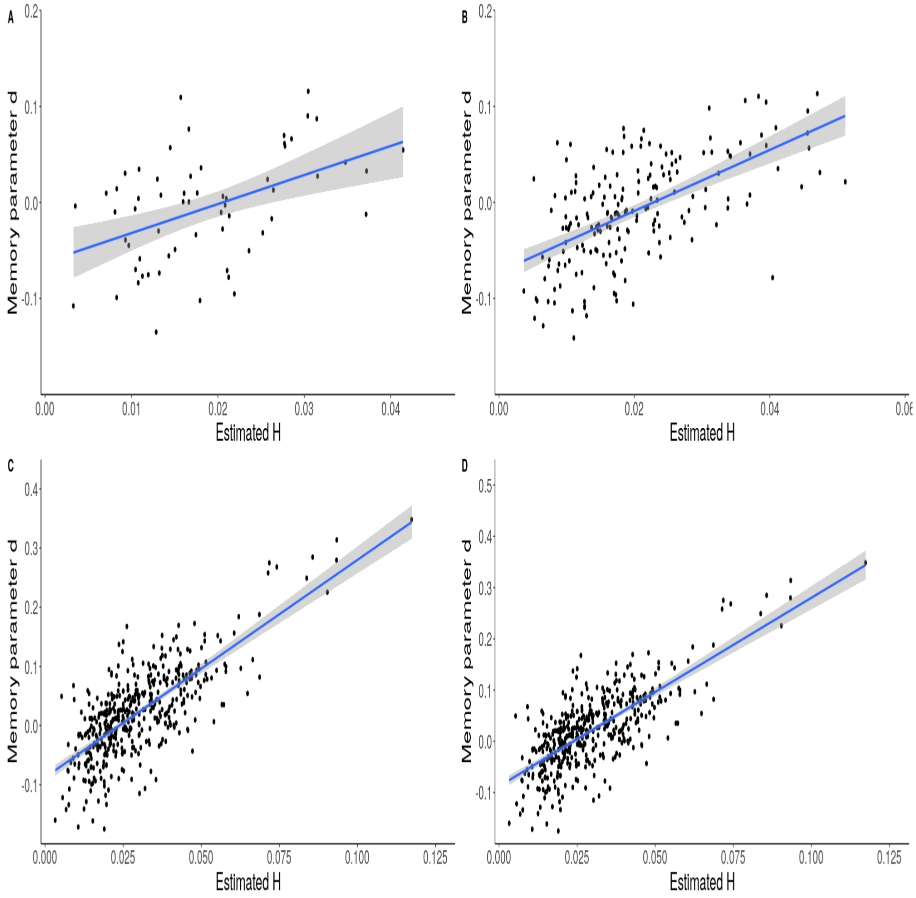


A. sub-17821 from ds000258; B. sub-21300 from ds000258, C. sub-28 from ds000210, D. sub-30 from ds000210. The number of datapoints is determined by the number of voxels inside the ventricles, which is determined by the size of the participant's ventricles. $d$ = ARFIMA$[0, d, 0]$ memory parameter, $H$ = Hurst parameter

Figure 14: Small air bubble in phantom 1 across the eight echoes

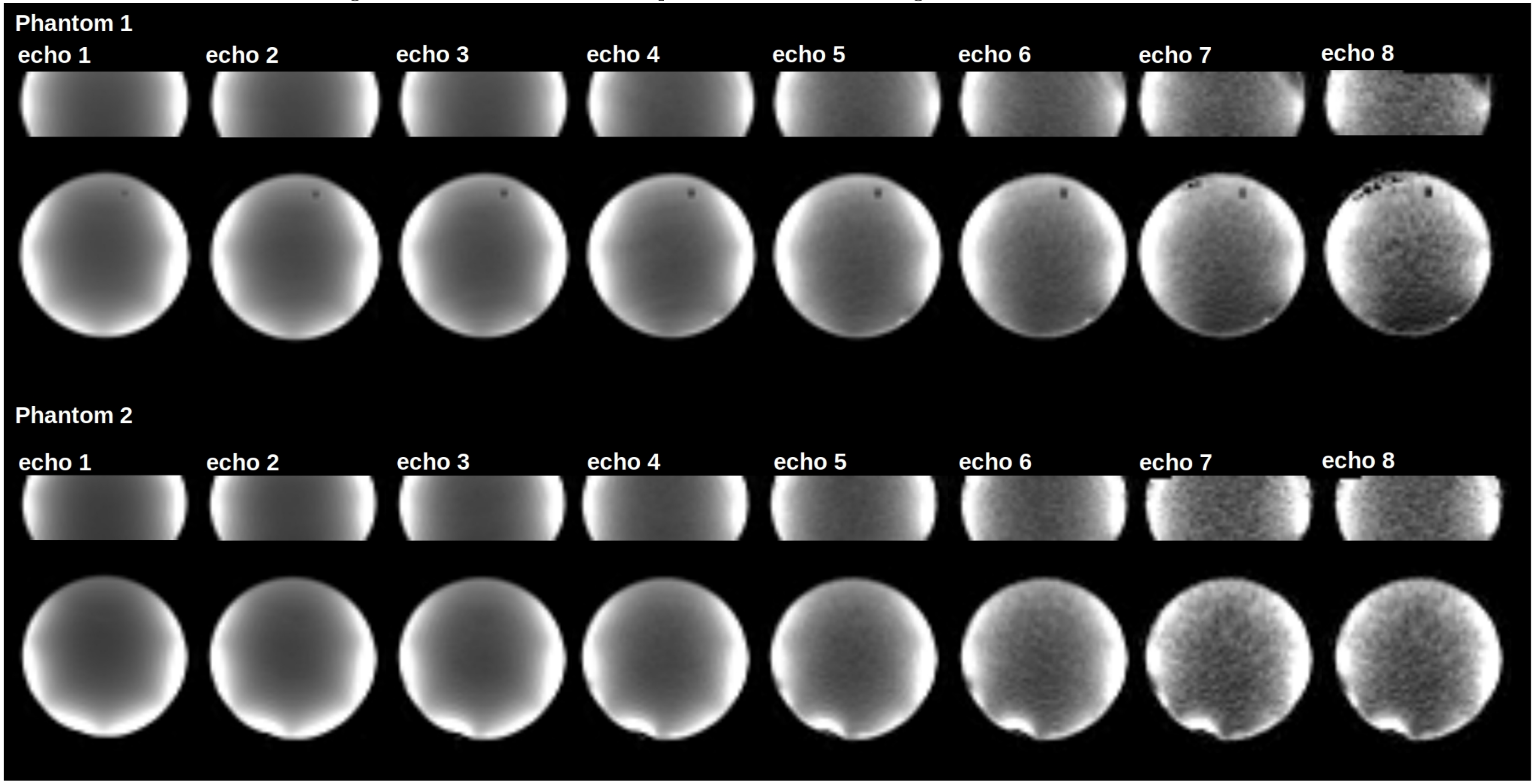
